# Cross-neutralization of a SARS-CoV-2 antibody to a functionally conserved site is mediated by avidity

**DOI:** 10.1101/2020.08.02.233536

**Authors:** Hejun Liu, Nicholas C. Wu, Meng Yuan, Sandhya Bangaru, Jonathan L. Torres, Tom G. Caniels, Jelle van Schooten, Xueyong Zhu, Chang-Chun D. Lee, Philip J.M. Brouwer, Marit J. van Gils, Rogier W. Sanders, Andrew B. Ward, Ian A. Wilson

**Affiliations:** Department of Integrative Structural and Computational Biology, The Scripps Research Institute, La Jolla, CA 92037, USA; Department of Medical Microbiology, Amsterdam UMC, University of Amsterdam; Department of Microbiology and Immunology, Weill Medical College of Cornell University, New York, NY 10021, USA; IAVI Neutralizing Antibody Center, The Scripps Research Institute, La Jolla, CA 92037, USA; Consortium for HIV/AIDS Vaccine Development (CHAVD), The Scripps Research Institute, La Jolla, CA 92037, USA; The Skaggs Institute for Chemical Biology, The Scripps Research Institute, La Jolla, CA, 92037, USA

## Abstract

Most antibodies isolated from COVID-19 patients are specific to SARS-CoV-2. COVA1-16 is a relatively rare antibody that also cross-neutralizes SARS-CoV. Here we determined a crystal structure of COVA1-16 Fab with the SARS-CoV-2 RBD, and a negative-stain EM reconstruction with the spike glycoprotein trimer, to elucidate the structural basis of its cross-reactivity. COVA1-16 binds a highly conserved epitope on the SARS-CoV-2 RBD, mainly through a long CDR H3, and competes with ACE2 binding due to steric hindrance rather than epitope overlap. COVA1-16 binds to a flexible up conformation of the RBD on the spike and relies on antibody avidity for neutralization. These findings, along with structural and functional rationale for the epitope conservation, provide a blueprint for development of more universal SARS-like coronavirus vaccines and therapies.

## MAIN

The ongoing coronavirus infectious disease 2019 (COVID-19) pandemic of severe acute respiratory syndrome coronavirus 2 (SARS-CoV-2) [1] is unlikely to end anytime soon [2]. Given the current lack of protective vaccines and antivirals, virus clearance and recovery of SARS-CoV-2 patients have to rely mainly on the generation of a neutralizing antibody response. To date, most neutralizing antibodies from convalescent patients target the receptor-binding domain (RBD) on the trimeric spike (S) glycoprotein [3–7], whose natural function is to mediate viral entry by first attaching to the human receptor angiotensin–converting enzyme 2 (ACE2) and then fusing its viral membrane with the host cell [1, 8–11]. SARS-CoV-2 is phylogenetically closely related to SARS-CoV [1], which caused the 2002-2003 human epidemic. However, SARS-CoV-2 and SARS-CoV only share 73% amino-acid sequence identity in their RBD, compared to 90% in their S2 fusion domain. Nevertheless, a highly conserved epitope on the SARS-CoV-2 RBD was previously identified from studies of a SARS-CoV neutralizing antibody CR3022 [12, 13], which was originally isolated almost 15 years ago [14]. Many human monoclonal antibodies have now been shown to target the SARS-CoV-2 S protein [3–7, 13, 15–24], but cross-neutralizing antibodies are relatively uncommon in COVID-19 patients [5, 6, 19, 25]. To date, the only structurally characterized cross-neutralizing human antibodies are S309 [18] and ADI-56046 [17] from SARS-CoV survivors, as well as EY6A from a COVID-19 patient [26]. Such structural and molecular characterization of cross-neutralizing antibodies is extremely valuable for therapeutic and vaccine design to confer broader protection against human SARS-like viruses that include the extensive reservoir of zoonotic coronaviruses in bats, camels, pangolins etc.

Antibody COVA1-16 was recently isolated from a convalescent COVID-19 patient and can cross-neutralize both SARS-CoV-2 (IC_50_ 0.13 µg/mL) and SARS-CoV (IC_50_ 2.5 µg/mL) pseudovirus [6]. The heavy and light chains of COVA1-16 are encoded by IGHV1-46, IGHD3-22, IGHJ1, and by IGKV1-33, IGKJ4, with a relatively long complementarity determining region (CDR) H3 of 20 amino acids (Figure S1). IGHV of COVA1-16 is only 1% somatically mutated at the nucleotide sequence level (one amino-acid change) from the germline gene, whereas its IGKV is 1.4% somatically mutated (three amino-acid changes). Here we determined the crystal structure of COVA1-16 in complex with SARS-CoV-2 RBD at 2.89 Å resolution to identify its binding site (epitope) and mechanism of cross-neutralization (Figure 1A, Table S1). The epitope of COVA1-16 overlaps extensively with that of CR3022, but also extends towards the periphery of the ACE2 binding site (Figure 1B) [13]. Seventeen out of 25 residues in the COVA1-16 epitope overlap with the CR3022 binding site (17 of 28 residues) (Figure 1C). Consistent with structural identification of its epitope, COVA1-16 can compete with CR3022 for RBD binding (Figure S2). COVA1-16 appears to have some resemblance to SARS-CoV cross-neutralizing antibody ADI-56046, whose epitope appears to span both the CR3022 epitope and ACE2-binding site, as indicated by negative-stain electron microscopy (nsEM) [17]. Interestingly, COVA1-16 also competes with ACE2 for RBD binding (Figure S2) [6], although its epitope does not overlap the ACE2 binding site (Figure 1B). Therefore, COVA1-16 inhibits ACE2 binding due to steric hindrance with its light chain rather than by direct interaction with the receptor binding site (Figure 1D).

**Figure 1.**
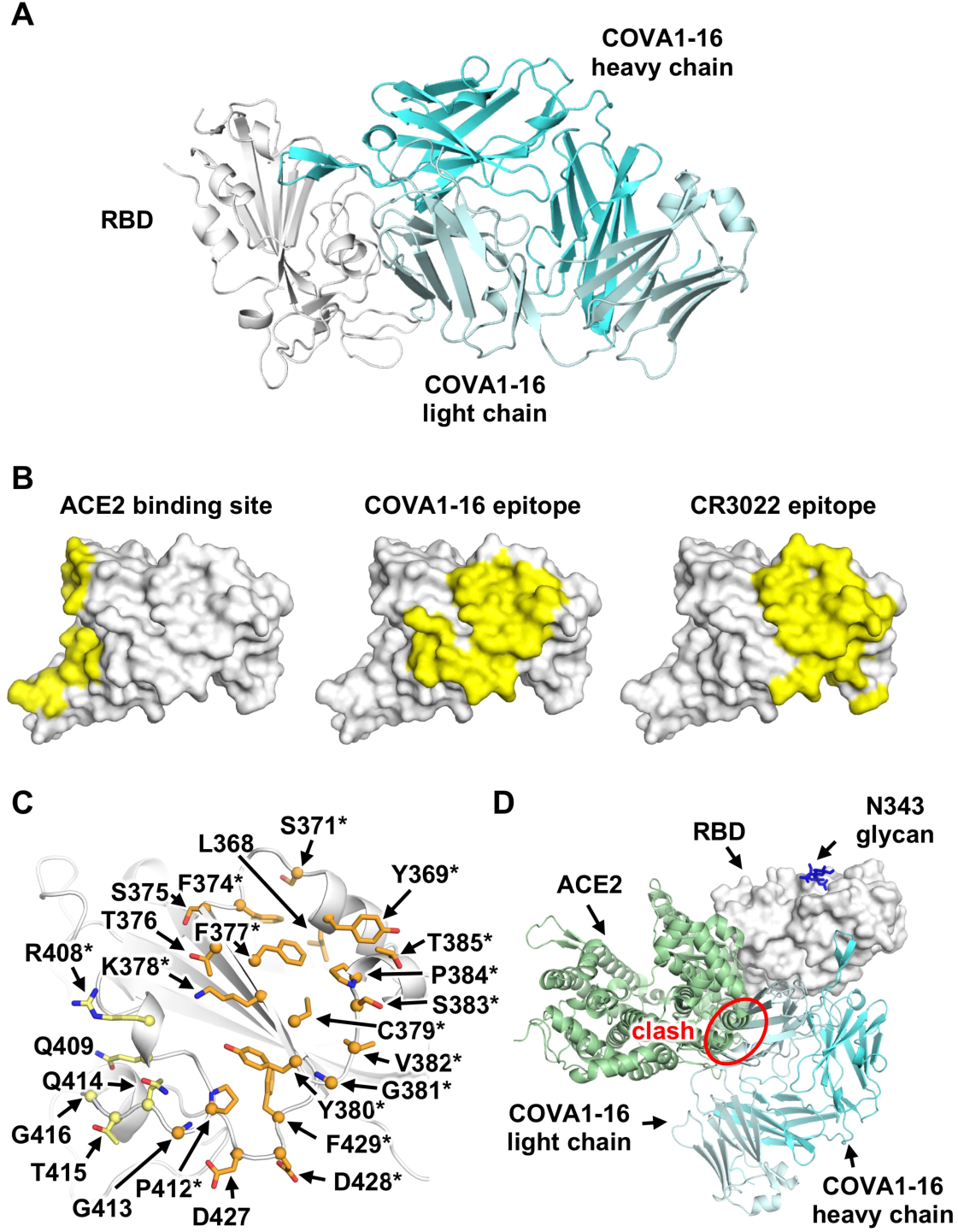
Comparison of COVA1-16 binding mode with CR3022 and ACE2. **(A)** Crystal structure of COVAI -16/RBD complex with RBD in grey and COVA1-16 Fab in cyan (heavy chain) and greyish blue (light chain). **(B)** ACE2-binding site (PDB 6M0J, left) [10], COVA1-16 epitope (this study, middle), and CR3022 epitope (PDB 6W41, right) [13] are highlighted in yellow. **(C)** RBD residues in the COVA1-16 epitope are shown. Epitope residues contacting the heavy chain are in orange and light chain in yellow. Representative epitope residues are labeled. Residues that are also part of CR3022 epitope are indicated with asterisks. **(D)** The ACE2/RBD complex structure is aligned in the same orientation as the COVA1-16/RBD complex. COVA1-16 (cyan) would clash with ACE2 (green) if they were to approach their respective RBD binding sites at the same time (indicated by red circle).

The RBD can adopt up and down conformations on the S trimer [27, 28]. While the ACE2 receptor only binds the RBD in the up conformation [9], previously characterized cross-neutralizing antibodies S309 from a convalescent SARS-CoV patient and COVA2-15 from a SARS-CoV-2 patient [6], can bind the RBD in both up and down conformations [18, 27]. However, unlike S309, the COVA1-16 epitope is completely buried when the RBD is in the down conformation (Figure 2A), akin to the CR3022 epitope [13]. Even in the up conformation of the RBD on an unliganded SARS-CoV-2 S trimer [27], the epitope of COVA1-16 would not be fully exposed (Figure 2A). We thus performed nsEM analysis of COVA1-16 in complex with the SARS-CoV-2 S trimer (Figure 2B). Three-dimensional (3D) reconstructions revealed that COVA1-16 can bind to a range of RBD orientations on the S protein when in the up position, indicating its rotational flexibility (Figure 2C). COVA1-16 can bind the S trimer either from the top (i.e. perpendicular to the trimer apex, Figure 2C, yellow, blue and pink) or from the side (i.e. more tilted, Figure 2C, brown). Model fitting of the COVA1-19/RBD crystal structure into the nsEM map indicates that the RBD on the S trimer is more open around the apex when COVA1-16 binds compared to unliganded trimers (Figure S3A-B). Bivalent binding of the COVA1-16 IgG between adjacent S trimers also appears to be plausible (Figure S3C). A recent cryo-electron tomography (cryo-ET) analysis demonstrated that the average distance between prefusion S on the viral surface is around 150 Å [29], which is comparable to the distance spanned between the tip of the two Fabs on an IgG (typically around 100 Å to 150 Å, although longer distances have been observed) [30]. Indeed, COVA1-16 IgG binds much more tightly than Fab to SARS-CoV-2 RBD, with dissociation constants (K_D_) of 0.2 nM and 46 nM, respectively (Figure S4A), reflecting bivalent binding in the assay format. Similarly, COVA1-16 IgG binds more strongly than Fab to SARS-CoV RBD (K_D_ of 125 nM vs 405 nM) (Figure S4B). Moreover, the apparent affinity of COVA1-16 IgG decreased to approximately the Fab value when the amount of SARS-CoV-2 RBD loaded on the biosensor was decreased, substantiating the notion that COVA1-16 can bind bivalently in this assay (Figure S4C).

**Figure 2.**
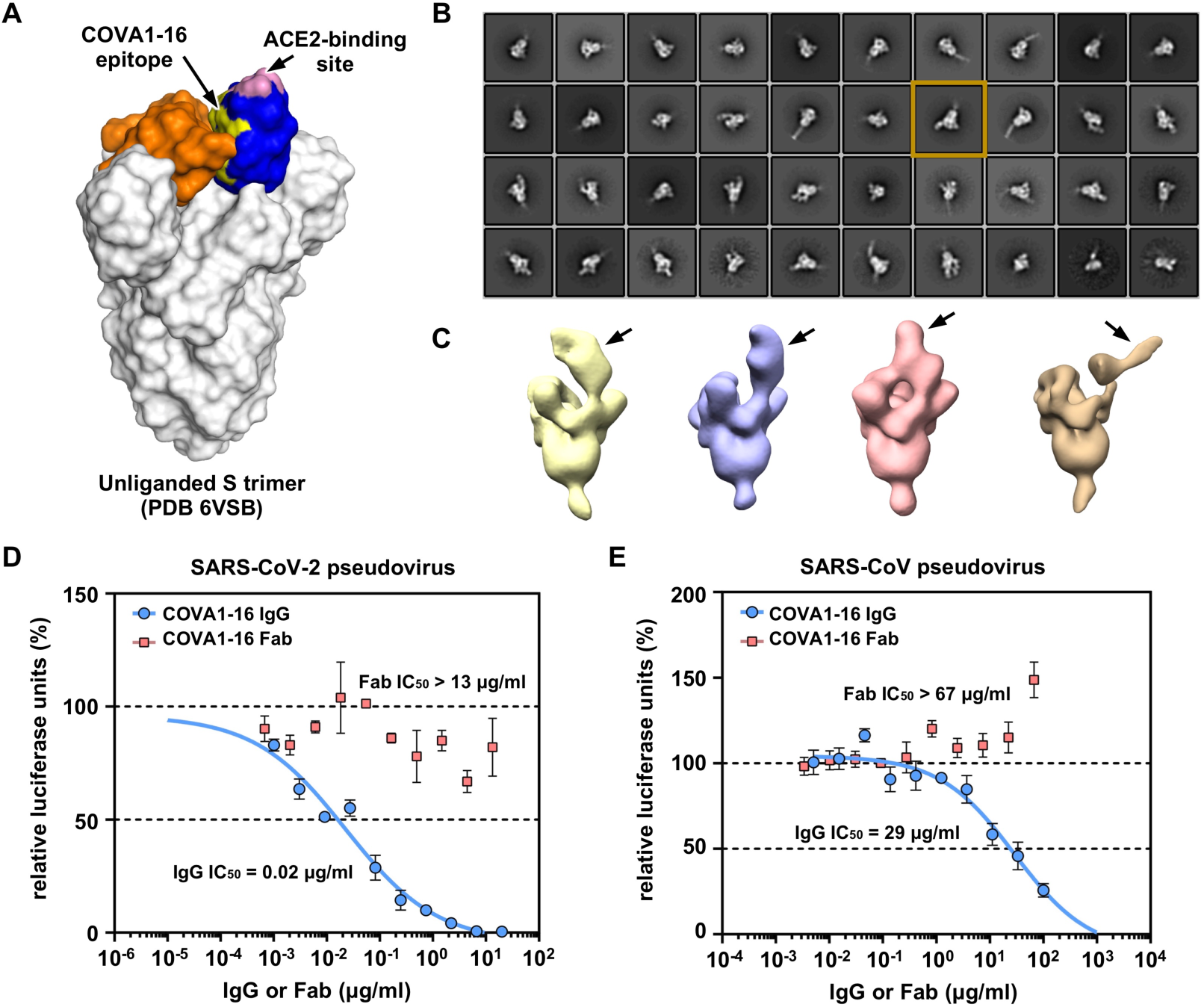
Negative-stain electron microscopy analysis and IgG avidity effect of COVA1-16. **(A)** The COVA1-16 epitope on the unliganded SARS-CoV-2 S trimer with one RBD in the “up” conformation (blue) and two in the “down” conformation (orange) (PDB 6VSB) [27]. COVA1-16 epitope is in yellow and ACE2-binding site in pink. **(B)** Representative 2D class averages from negative-stain EM analysis of SARS-CoV-2 S trimer complexed with COVA1-16 Fab. The 2D class corresponding to the most outward conformation of COVA-16 Fab in complex with S trimer is highlighted in a mustard box. **(C)** Various conformations of COVA1-16 Fab in complex with the S trimer is revealed by 3D reconstructions. The location of COVA1-16 Fab is indicated by an arrow. **(D-E)** Neutralization activities of COVA1-16 IgG (blue) and Fab (red) against **(D)** SARS-CoV-2 and **(E)** SARS-CoV are measured in a luciferase-based pseudovirus assay. The half maximal inhibitory concentrations (IC_50_s) for IgG and Fab are indicated in parenthesis. Of note, neutralization for the IgG (IC_50_ = 0.08 µg/mL) against SARS-CoV-2 pseudovirus infecting 293T/ACE2 cells is comparable to that measured in Huh7 cells (IC_50_ = 0.13 µg/mL) as reported previously [6].

Bivalent IgG binding is also important for the neutralization activity of COVA1-16 (Figure 2D-E). COVA1-16 IgG neutralizes SARS-CoV-2 pseudovirus with a half maximal inhibitory concentration (IC_50_) of 0.02 µg/mL, which is similar to that previously measured for SARS-CoV-2 pseudovirus (IC_50_ of 0.13 µg/mL) [6]. In contrast, COVA1-16 Fab does not neutralize SARS-CoV-2 pseudovirus even up to 13 µg/mL. A similar effect is also observed for SARS-CoV pseudovirus, which is neutralized by COVA1-16 IgG at an IC_50_ of 29 µg/mL, but not by COVA1-16 Fab even up to 67 µg/mL (Figure 2E). Of note, COVA1-16 is less potent against authentic SARS-CoV-2 (IC_50_ = 0.75 µg/mL) [6]. Whether such a difference is due to variation in S protein density on the viral surface versus pseudovirus or to other factors deserves future investigation. It will also be informative to compare the number, density and conformational states of the S proteins on SARS-CoV-2 and SARS-CoV virions. Overall, our findings support the importance of bivalent binding for SARS-CoV-2 neutralizing antibodies, and especially for cross-neutralization of SARS-CoV. Such a contribution of bivalent IgG (avidity) to SARS-CoV-2 neutralization has also been suggested in a recent study that compared binding of polyclonal IgGs and Fabs [24]. Furthermore, a single-domain camelid antibody VHH-72 dramatically improved its neutralization activity to SARS-CoV-2 when expressed as a bivalent Fc-fusion [31]. These observations are similar to some influenza broadly neutralizing antibodies to the hemagglutinin (HA) receptor binding site, where bivalent binding can increase avidity and neutralization breadth [32, 33].

Next we examined the molecular details of the interactions between COVA1-16 and SARS-CoV-2. COVA1-16 binding to the RBD is dominated by the heavy chain, which accounts for 82% of its total buried surface area (BSA, 694 Å^2^ out of a total of 844 Å^2^). Most of the interactions are mediated by CDR H3 (Figure 3A), which contributes 70% (594 Å^2^) of the total BSA. CDR H3 forms a beta-hairpin with a type I beta-turn at its tip and is largely encoded by IGHD3-22 (from V_H_ N98 to V_H_ Y100f, Figure S1C and Figure 3B). The beta-hairpin conformation is stabilized by four main chain-main chain hydrogen-bonds (H-bonds) and a side chain-side chain H-bond between V_H_ N98 and V_H_ Y100f at either end of the IGHD3-22-encoded region (Figure 3B). Four H-bonds between the tip of CDR H3 and the RBD are formed from two main chain-main chain interactions with RBD C379, and two with V_H_ R100b (Table S2). The positively charged guanidinium of V_H_ R100b also interacts with the negative dipole at the C-terminus of a short α-helix in the RBD (residues Y365 to Y369). Interestingly, V_H_ R100b is a somatically mutated residue (codon = AGG in the IGHD3-22-encoded region, where the germline residue is a Ser (codon = AGT, Figure S1C). The short Ser side chain would likely not contact the RBD nor provide electrostatic complementarity. Interestingly, a somatic revertant V_H_ R100bS actually improved binding affinity of COVA1-16 to the RBD, mostly due to an increased on-rate (Figure S5). Nevertheless, COVA1-16 has a much slower off-rate than its V_H_ R100bS mutant, which may have led to its selection. The CDR H3 tip also interacts with the RBD through hydrophobic interactions between V_H_ Y99 and the aliphatic portion of RBD K378, as well as a π-π interaction between V_H_ Y100 and the RBD V382-S383 peptide backbone (Figure 3B). CDR H3 forms an additional four H-bonds with the RBD, involving the side chains of V_H_ R97 and Q101 (Figure 3B). We further determined the unliganded structure of COVA1-16 Fab to 2.53 Å resolution and found that the CDR H3 distal region was not resolved due to lack of electron density indicating its inherent flexibility (Figure S6). CDR H1 and CDR L2 of COVA1-16 also interact with the RBD, but much less so compared to CDR H3. The V_H_ T28 main chain and V_H_ Y32 side chain in CDR H1 H-bond with D427 (Figure 3C, Table S2), whereas V_L_ N53 in CDR L2 H-bonds with RBD R408 (Figure 3D, Table S2).

**Figure 3.**
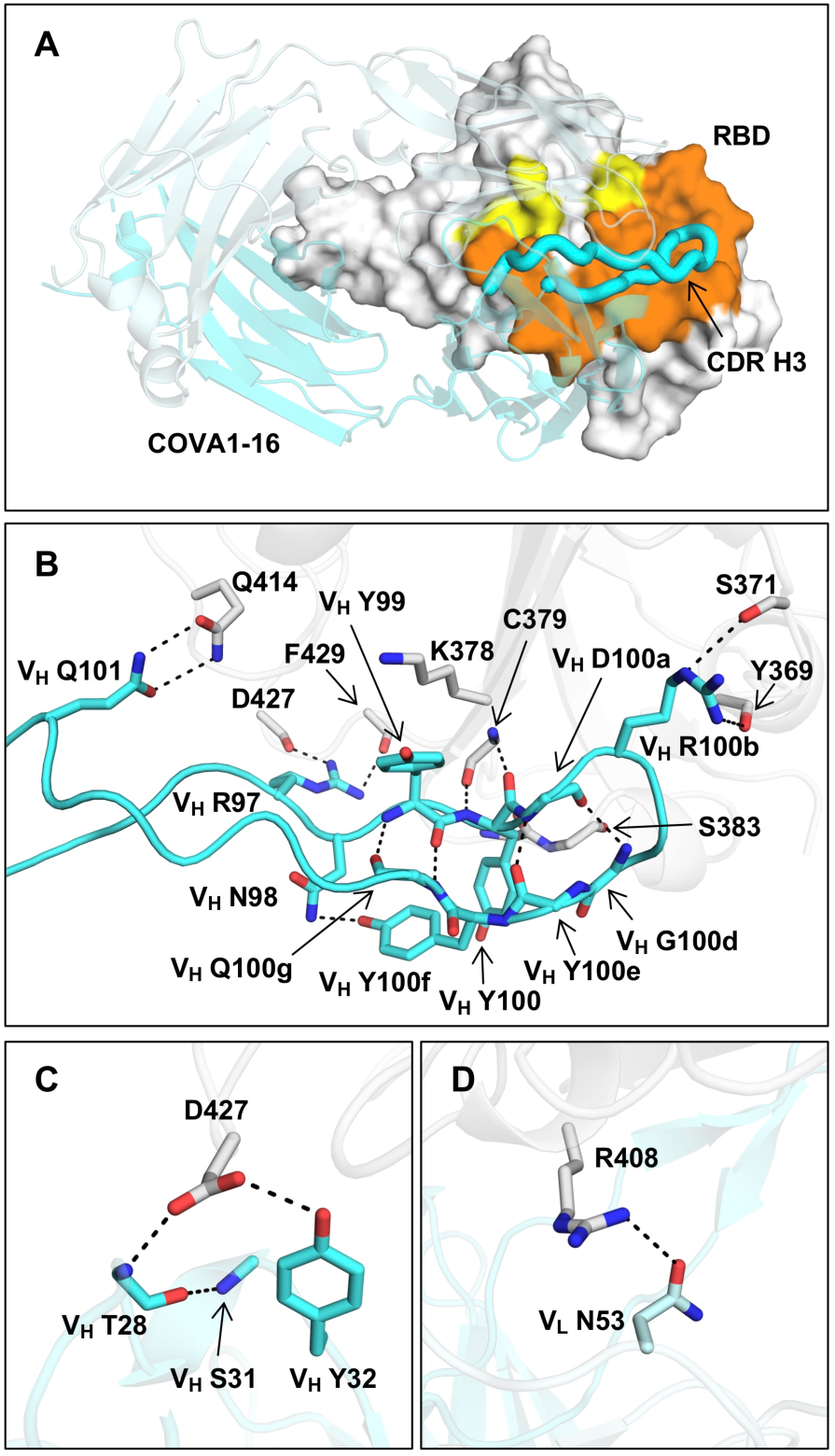
Interaction between SARS-CoV-2 RBD and COVA1-16. **(A)** The epitope of COVA1-16 is highlighted in yellow and orange. Epitope residues that are in contact with CDR H3 are in orange, and yellow otherwise. COVA1-16 (cyan) is in cartoon representation with CDR H3 depicted in a thick tube. The RBD (white) is in a surface representation. The BSA on COVA1-16 and RBD are 844 Å^2^ and 779 Å^2^, respectively. **(B)** Interactions of SARS-CoV-2 RBD (white) with **(B)** CDR H3, **(C)** CDR H1, and **(D)** CDR L2 of COVA1-16 (cyan) are shown. Hydrogen bonds are represented by dashed lines. In **(C)**, a 3_10_ turn is observed in CDR H1 for residues V_H_ T28 to V_H_ S31.

CDR H3-dominant antibodies have been seen in the human immune response to other viral pathogens. Striking examples are antibodies PG9 and PG16, whose CDR H3s interact extensively along their length with the apex of the HIV-1 Envelope protein [34, 35]. Another example is C05, which is essentially a single loop binder that inserts its very long CDR H3 (24 residues) into the RBD of influenza HA [32], thereby providing a template for design of a high-avidity protein inhibitor of influenza virus, where the H3 loop was fused to a scaffold protein [36]. The long CDR H3 of COVA1-16 may similarly facilitate therapeutic designs that could also include peptide-based antivirals, as exemplified by a potent cyclic peptide fusion inhibitor of influenza HA [37, 38].

Compared to the ACE2-binding site, the COVA1-16 epitope is much more highly conserved among SARS-CoV-2, SARS-CoV, and other SARS-related coronaviruses (SARSr-CoV) (Figure 4A-D, Figure S7 and Figure S8) [6]. To investigate possible structural and functional reasons for this sequence conservation, we analyzed the epitope location in the context of the SARS-CoV-2 trimeric S protein with all RBDs in the “down” conformation [39] (Figure 4E and Figure S7). The COVA1-16 epitope is completely buried at the center of the trimer in the interface between the S1 and S2 domains and is largely hydrophilic (Figure S9). The polar side chains of K378, Q414, R408, and D427, which are involved in binding to COVA1-16, are all very close to the interface with adjacent protomers in the S trimer. Interestingly, the R408 side chain, which is positioned by Q414 via a H-bond, points towards a region in the adjacent protomer 2 with a positive electrostatic potential. Similarly, D427 is juxtaposed to a region in protomer 2 with a negative electrostatic potential. These repulsive charges would help favor the metastability required for transient opening and closing of the RBD in “up” and “down” conformations prior to ACE2 receptor binding. In contrast, the K378 side chain points towards a region in protomer 3 with negative electrostatic potential, thus favoring the “down” RBD conformation. Furthermore, in the “down” conformation, part of the COVA1-16 epitope interacts with the long helices formed from the heptad repeat motifs of S2 fusion domain (Figure 4E–F). Notably, S383 and T385 in the COVA1-16 epitope make three H-bonds with the tops of the helices and their connecting regions (Figure 4F). This mixture of attractive and repulsive forces would seem to be important for control of the dynamics of the RBD and, hence, for the biological function of the metastable pre-fusion S protein in receptor binding and fusion. The complementarity of fit of the epitope interface with the other RBDs and the S2 domain in the S trimer further explains the epitope conservation (Figure S10). Therefore, the high sequence conservation of the COVA1-16 epitope appears related to the functional requirement for this component of the RBD surface to be deeply buried within the S trimer in the “down” conformation.

**Figure 4.**
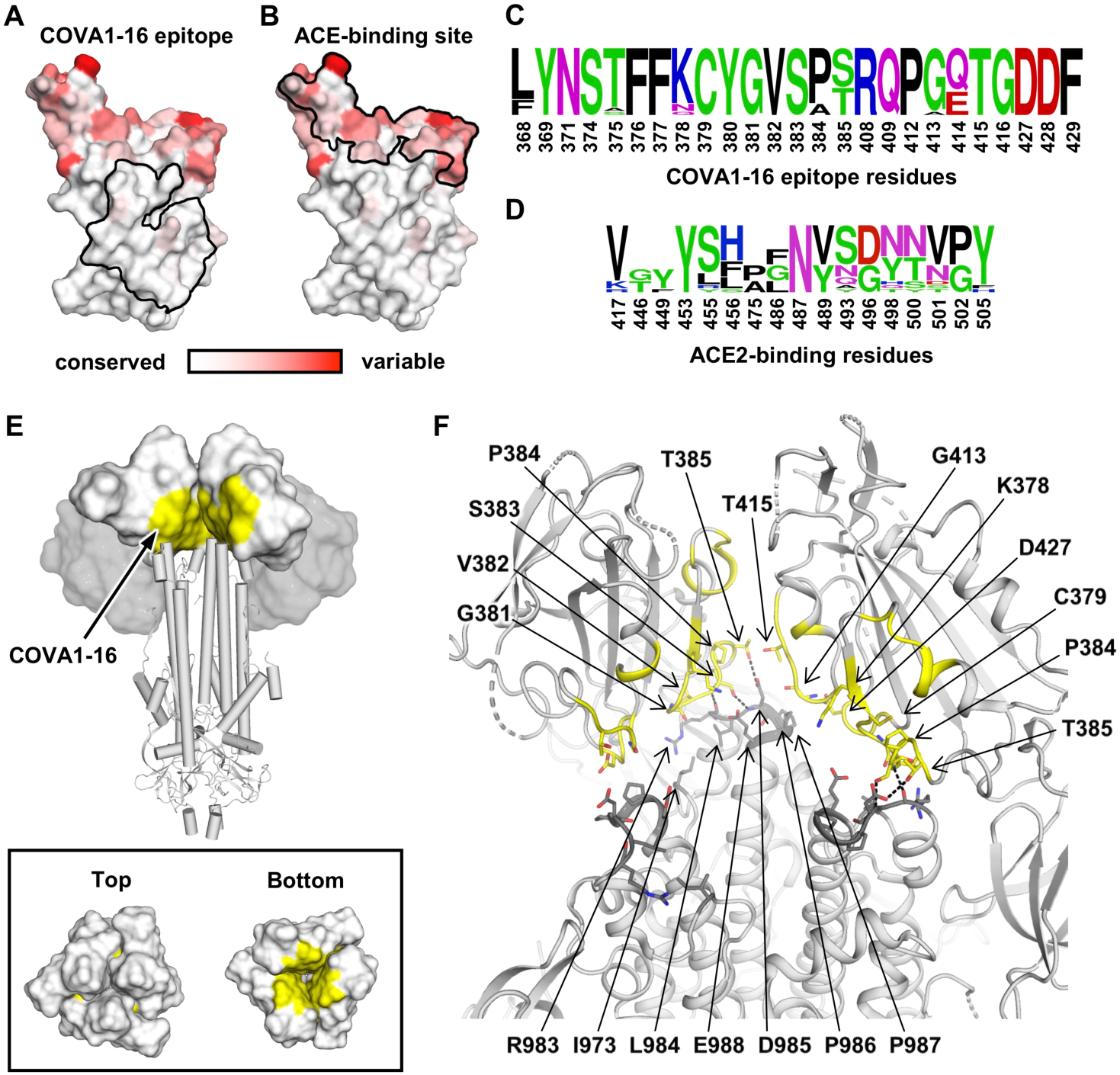
Sequence conservation of COVA1-16 epitope and ACE2-binding site. **(A–B)** Sequence conservation of the RBD among 17 SARS-like CoVs (Figure S7) is highlighted on the RBD structure with the **(A)** COVA1-16 epitope and **(B)** ACE2-binding site indicated by the black outline. The backside of this view is shown in Figure S8. **(C-D)** Sequence conservation of **(C)** COVA1-16 epitope and **(D)** ACE2-binding site is shown as a sequence logo. **(E)** Location of COVA1-16 epitope (yellow) on the SARS-CoV-2 S trimer when all three RBDs are in the down conformation (PDB 6VXX) [39]. RBDs are represented as a white surface, N-terminal domains (NTDs) as a grey surface, and the S2 domain in a cartoon representation. Top panel: for visualization of the COVA1-16 epitope, the RBD and NTD from one of the three protomers was removed. Bottom panel: top and bottom views of the COVA1-16 epitopes on the three RBDs in the “down” conformation. **(F)** COVA1-16 epitope is shown in yellow on a ribbon representation of a SARS-CoV-2 S trimer (PDB 6VXX) [39]. Epitope residues in the RBD involved in interaction with the S2 domain are shown in yellow sticks, and S2 domain interacting residues in dark grey sticks. Dashed lines indicate hydrogen bonds. Interface residues are calculated using PISA [54]. The S1 segment from the third protomer is omitted to clarify the view of the interfaces that the COVA1-16 epitope makes with the S2 domain.

From the SARS-CoV-2 RBD/antibody complex structures to date, a significant portion of the RBD surface can be targeted by antibodies (Figure 5). One surface not yet observed to be targeted is partially covered by N-glycans at residues N165 on the N-terminal domain (NTD) and N343 on the RBD, which may hinder B cell receptor access and create a “silent face” (Figure S11), although the N343 glycan is incorporated in the S309 epitope [18]. While antibodies that target the ACE2-binding site, such as BD23 [7], CB6 [23], B38 [20], P2B-2F6 [19], CC12.1 [40], and CC12.3 [40], do not show cross-neutralization activity to SARS-CoV, the conserved epitopes further from the ACE2-binding site seem to be more able to support cross-neutralization [13, 18, 26]. It is also interesting that these so far rare cross-neutralizing antibodies, including COVA1-16, often seem to bind to epitopes that are not readily accessible in the pre-fusion native structure [17, 26]. This finding is similar to a recent discovery in influenza virus, where a class of cross-protective antibodies target a conserved epitope in the trimeric interface of the HA [41–43]. Due to the inaccessibility of the COVA1-16 epitope on the S protein, it is possible that an RBD-based rather than S-based immunogen can elicit larger numbers of COVA1-16-like antibodies. Cross–neutralizing antibodies have also provided important insights into therapeutic and vaccine design, as for influenza virus [44] and HIV [45]. As SARS-CoV-2 continues to circulate in the human population and other zoonotic coronaviruses constitute future pandemic threats [46], it is certainly worth considering the development of more universal coronavirus vaccines and therapeutics that can cross-neutralize antigenically drifted SARS-CoV-2 viruses, as well as zoonotic SARS-like coronaviruses.

**Figure 5.**
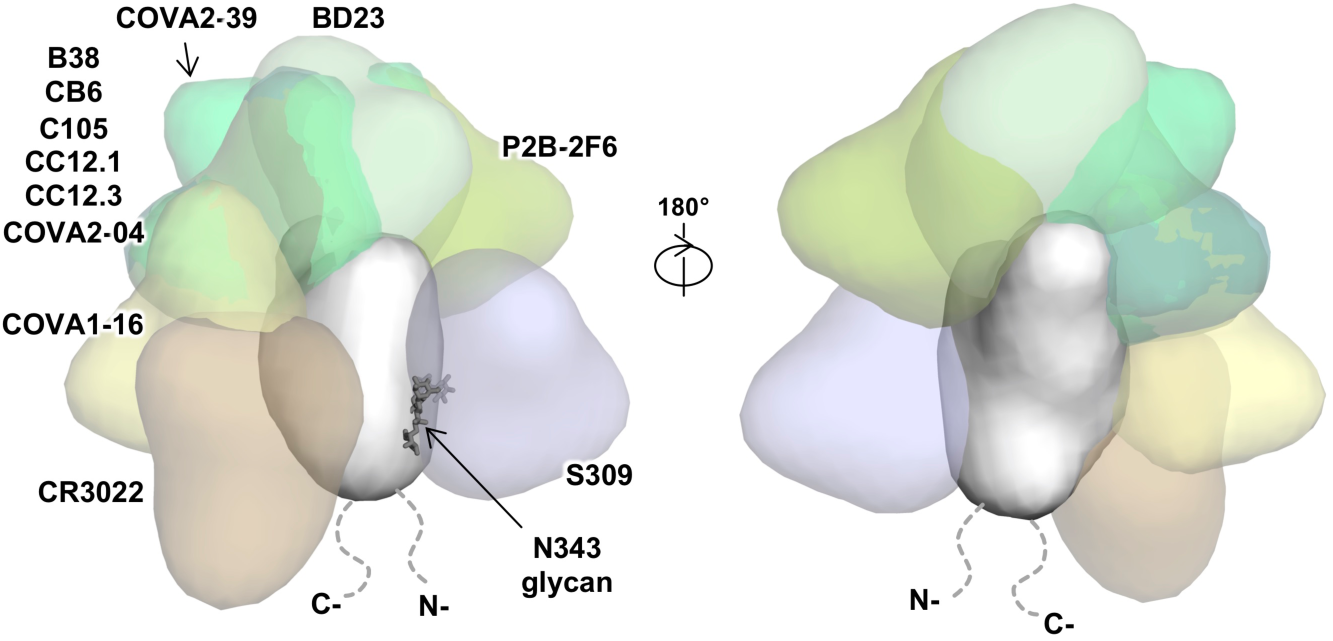
Interaction between SARS-CoV-2 RBD and structurally characterized antibodies. The binding of known SARS-CoV-2 RBD-targeting antibodies to the RBD is compared. The ACE2-binding site overlaps with epitopes of B38 (PDB 7BZ5) [20], C105 (6XCM) [24], CB6 (7C01) [23], CC12.1 (6XC3) [40], CC12.3 (6XC4) [40], BD23 (7BYR) [7], and P2B-2F6 (7BWJ) [19], but not the epitopes of COVA1-16 (this study), CR3022 (PDB 6W41) [13], COVA2-04 [63], COVA2-39 [63], and S309 (PDB 6WPS) [18]. Of note, while CR3022 only neutralizes SARS-CoV but not SARS-CoV-2 in *in vitro* assays [13], a recent study isolated an antibody (EY6A) that binds to a similar epitope as CR3022 and cross-neutralizes SARS-CoV-2 and SARS-CoV [26].

## MATERIALS AND METHODS

### Expression and purification of SARS-CoV-2 RBD

The receptor-binding domain (RBD) (residues 319-541) of the SARS-CoV-2 spike (S) protein (GenBank: QHD43416.1), and the RBD (residues 306-527) of the SARS-CoV S protein (GenBank: ABF65836.1), were cloned into a customized pFastBac vector [47], and fused with an N-terminal gp67 signal peptide and C-terminal His_6_ tag [13]. For each RBD, we further cloned a construct with an AviTag inserted in front of the His_6_ tag. To express the RBD, a recombinant bacmid DNA was generated using the Bac-to-Bac system (Life Technologies). Baculovirus was generated by transfecting purified bacmid DNA into Sf9 cells using FuGENE HD (Promega), and subsequently used to infect suspension cultures of High Five cells (Life Technologies) at an MOI of 5 to 10. Infected High Five cells were incubated at 28 °C with shaking at 110 r.p.m. for 72 h for protein expression. The supernatant was then concentrated using a 10 kDa MW cutoff Centramate cassette (Pall Corporation). The RBD protein was purified by Ni-NTA, followed by size exclusion chromatography, and buffer exchanged into 20 mM Tris-HCl pH 7.4 and 150 mM NaCl. For binding experiments, RBD with AviTag was biotinylated as described previously [32] and purified by size exclusion chromatography on a Hiload 16/90 Superdex 200 column (GE Healthcare) in 20 mM Tris-HCl pH 7.4 and 150 mM NaCl.

### Expression and purification of Fabs

Expression plasmids encoding the heavy and light chains of the COVA1-16 Fab were transiently co-transfected into ExpiCHO cells at a ratio of 2:1 (HC:LC) using ExpiFectamine™ CHO Reagent (Thermo Fisher Scientific) according to the manufacturer’s instructions. The supernatant was collected at 10 days post-transfection. The Fabs were purified with a CaptureSelect™ CH1-XL Affinity Matrix (Thermo Fisher Scientific) followed by size exclusion chromatography.

### Expression and purification of ACE2

The N-terminal peptidase domain of human ACE2 (residues 19 to 615, GenBank: BAB40370.1) was cloned into phCMV3 vector and fused with a C-terminal Fc tag. The plasmids were transiently transfected into Expi293F cells using ExpiFectamine™ 293 Reagent (Thermo Fisher Scientific) according to the manufacturer’s instructions. The supernatant was collected at 7 days post-transfection. Fc-tagged ACE2 protein was then purified with a Protein A column (GE Healthcare) followed by size exclusion chromatography.

### Crystallization and x-ray structure determination

The COVA1-16 Fab complex with RBD was formed by mixing each of the protein components in an equimolar ratio and incubating overnight at 4°C. The COVA1-16 Fab/RBD complex and COVA1-16 Fab apo (unliganded) protein were adjusted to around 11 mg/mL and screened for crystallization using the 384 conditions of the JCSG Core Suite (Qiagen) on our custom-designed robotic CrystalMation system (Rigaku) at Scripps Research. Crystallization trials were set-up by the vapor diffusion method in sitting drops containing 0.1 µl of protein and 0.1 µl of reservoir solution. Crystals used for x-ray data collection were harvested from drops containing 0.2 M sodium iodide and 20% (w/v) polyethylene glycol 3350 for the COVA1-16 Fab/RBD complex and from drops containing 0.08 M acetate pH 4.6, 20% (w/v) polyethylene glycol 4000, 0.16 M ammonium sulfate and 20% (v/v) glycerol for the COVA1-16 Fab. Crystals appeared on day 3, were harvested on day 7, pre-equilibrated in cryoprotectant containing 20% glycerol, and then flash cooled and stored in liquid nitrogen until data collection. Diffraction data were collected at cryogenic temperature (100 K) at Stanford Synchrotron Radiation Lightsource (SSRL) on the new Scripps/Stanford beamline 12-1 with a beam wavelength of 0.97946 Å, and processed with HKL2000 [48]. Structures were solved by molecular replacement using PHASER [49]. The models for molecular replacement of RBD and COVA1-16 were from PDB 6XC4 [40], 4IMK [50] and 2Q20 [51]. Iterative model building and refinement were carried out in COOT [52] and PHENIX [53], respectively. Epitope and paratope residues, as well as their interactions, were identified by accessing PISA at the European Bioinformatics Institute (http://www.ebi.ac.uk/pdbe/protint/pistart.html) [54].

### Expression and purification of recombinant S proteins

The SARS-CoV-2 S construct used for negative stain EM contains the mammalian-codon-optimized gene encoding residues 1-1208 of the S protein (GenBank: QHD43416.1), followed by a C-terminal T4 fibritin trimerization domain, an HRV3C cleavage site, 8x-His tag and a Twin-strep tags subcloned into the eukaryotic-expression vector pcDNA3.4. Three amino-acid mutations were introduced into the S1/S2 cleavage site (RRAR to GSAS) to prevent cleavage and two stabilizing proline mutations (K986P and V987P) to the HR1 domain. For additional S stabilization, residues T883 and V705 were mutated to cysteines to introduce a disulphide bond. The S plasmid was transfected into 293F cells and supernatant was harvested at 6 days post transfection. S protein was purified by running the supernatant through a streptactin column and then by size exclusion chromatography using a Superose 6 increase 10/300 column (GE Healthcare Biosciences). Protein fractions corresponding to the trimeric S protein were collected and concentrated.

### ns-EM sample preparation and data collection

SARS-COV-2 S protein was complexed with 3x molar excess of Fab at 30 minutes prior to direct deposition onto carbon-coated 400-mesh copper grids. The grids were stained with 2 % (w/v) uranyl-formate for 90 seconds immediately following sample application. Grids were either imaged at 200 KeV or at 120 KeV on a Tecnai T12 Spirit using a 4kx4k Eagle CCD. Micrographs were collected using Leginon [55] and the images were transferred to Appion for processing. Particle stacks were generated in Appion [56] with particles picked using a difference-of-Gaussians picker (DoG-picker) [57]. Particle stacks were then transferred to Relion [58] for 2D classification followed by 3D classification to sort well-behaved classes. Selected 3D classes were auto-refined on Relion and used to make figures with UCSF Chimera.

### Protein expression and purification for antibody binding studies

All constructs were expressed transiently in HEK293F (Invitrogen, cat no. R79009) cells maintained in Freestyle medium (Life Technologies). For soluble RBD proteins, cells were transfected at a density of 0.8-1.2 million cells/mL by addition of a mix of PEImax (1 µg/µL) with expression plasmids (312.5 µg/L) in a 3:1 ratio in OptiMEM. Supernatants of the soluble RBD proteins were harvested six days post transfection, centrifuged for 30 min at 4000 rpm and filtered using 0.22 µm Steritop filters (Merck Millipore). Constructs with a His_6_-tag were purified by affinity purification using Ni-NTA agarose beads. Protein eluates were concentrated, and buffer exchanged to PBS using Vivaspin filters with a 10 kDa molecular weight cutoff (GE Healthcare). Protein concentrations were determined by Nanodrop using the proteins peptidic molecular weight and extinction coefficient as determined by the online ExPASy software (ProtParam). For the COVA1-16 IgG 1 antibody, suspension HEK293F cells (Invitrogen, cat no. R79007) were cultured in FreeStyle medium (Gibco) and co-transfected with the two IgG plasmids expressing the corresponding HC and LC in a 1:1 ratio at a density of 0.8-1.2 million cells/mL in a 1:3 ratio with 1 mg/L PEImax (Polysciences). The recombinant IgG antibodies were isolated from the cell supernatant after five days as described previously (20, 48). In short, the cell suspension was centrifuged 25 min at 4000 rpm, and the supernatant was filtered using 0.22 µm pore size SteriTop filters (Millipore). The filtered supernatant was run over a 10 mL protein A/G column (Pierce) followed by two column volumes of PBS wash. The antibodies were eluted with 0.1 M glycine pH 2.5, into the neutralization buffer of 1 M TRIS pH 8.7 in a 1:9 ratio. The purified antibodies were buffer exchanged to PBS using 100 kDa VivaSpin20 columns (Sartorius). The IgG concentration was determined on the NanoDrop 2000 and the antibodies were stored at 4°C until further analyses.

### Measurement of binding affinities using biolayer interferometry

To determine the binding affinity of COVA1-16 IgG and His-tagged Fabs, 20 µg/mL of His-tagged SARS-CoV or SARS-CoV-2 RBD protein in running buffer (PBS, 0.02% Tween-20, 0.1% BSA) was loaded on Ni-NTA biosensors (ForteBio) for 300 s. Streptavidin biosensors (ForteBio) were used if the RBD was biotinylated. Next, the biosensors were transferred to running buffer containing IgG or Fab to determine the association rate, after which the sensor was transferred to a well containing running buffer to allow dissociation. As negative control, an anti-HIV-1 His-tagged Fab was tested at the highest concentration used for COVA1-16 Fab (400 nM). After each cycle, the sensors were regenerated by alternating 20 mM glycine in PBS and running buffer three times, followed by reactivation in 20 mM NiCl_2_ for 120 s. All steps were performed at 1000 rpm shaking speed. K_D_S were determined using ForteBio Octet CFR software. The avidity effects of IgG were investigated by titrating the SARS-CoV-2 RBD concentration (5, 1, 0.2 and 0.04 µg/mL) followed by loading on Ni-NTA biosensors for 480 s with an additional loading step with His-tagged HIV-1 gp41 for 480 s to minimize background binding of His-tagged Fabs to the biosensor. All other steps were performed as described above.

### Competition studies of antibodies with ACE-2 receptor

For competition assays, COVA1-16 IgG, CR3022 IgG, and human ACE2-Fc were all diluted to 250 nM. Ni-NTA biosensors were used. In brief, the assay has five steps: 1) baseline: 60 s with 1x kinetics buffer; 2) loading: 180 s with 20 µg/mL, His_6_-tagged SARS-CoV-2 RBD proteins; 3) baseline: 150 s with 1x kinetics buffer; 4) first association: 300 s with CR3022 IgG or human ACE2-Fc; and 5) second association: 300 s with human ACE2-Fc, CR3022 IgG, or COVA1-16 IgG.

### Pseudovirus neutralization assay

Neutralization assays were performed using SARS-CoV and SARS-CoV-2 S-pseudotyped HIV-1 virus and HEK-293T/ACE2 cells as described previously [59]. In brief, pseudotyped virus was produced by co-transfecting expression plasmids of SARS-CoV S and SARS-COV-2_Δ19_ S proteins (GenBank; AAP33697.1 and MT449663.1, respectively) with an HIV backbone expressing NanoLuc luciferase (pHIV-1_NL4-3_ ΔEnv-NanoLuc) in HEK293T cells (ATCC, CRL-11268). After 3 days, the cell culture supernatants containing SARS-CoV and SARS-CoV-2 S-pseudotyped HIV-1 viruses were stored at −80°C. HEK-293T/ACE2 cells were seeded 10,000 cells/well in a 96-well plate one day prior to the start of the neutralization assay. To determine the neutralizing capacity of COVA1-16 IgG and His_6_-tagged Fab, 20 or 100 µg/mL COVA1-16 IgG and equal molar of COVA1-16 Fab were serially diluted in 3-fold steps and mixed with SARS-CoV or SARS-CoV-2 pseudotyped virus and incubated for 1 h at 37°C. The pseudotyped virus and COVA1-16 IgG/Fab mix were then added to the HEK-293T/ACE2 cells and incubated at 37°C. After 48 h, cells were washed twice with PBS (Dulbecco’s Phosphate-Buffered Saline, eBiosciences) and lysis buffer was added. Luciferase activity of cell lysate was measured using the Nano-Glo Luciferase Assay System (Promega) and GloMax Discover System. The inhibitory concentration (IC_50_) was determined as the concentration of IgG or Fab that neutralized 50% of the pseudotyped virus using GraphPad Prism software (version 8.3.0).

### Sequence conservation analysis

RBD protein sequences from SARS-CoV and SARS-related coronavirus (SARSr-CoV) strains were retrieved from the following accession codes:

- GenBank ABF65836.1 (SARS-CoV)
- GenBank ALK02457.1 (Bat SARSr-CoV WIV16)
- GenBank AGZ48828.1 (Bat SARSr-CoV WIV1)
- GenBank ACU31032.1 (Bat SARSr-CoV Rs672)
- GenBank AIA62320.1 (Bat SARSr-CoV GX2013)
- GenBank AAZ67052.1 (Bat SARSr-CoV Rp3)
- GenBank AIA62300.1 (Bat SARSr-CoV SX2013)
- GenBank ABD75323.1 (Bat SARSr-CoV Rf1)
- GenBank AIA62310.1 (Bat SARSr-CoV HuB2013)
- GenBank AAY88866.1 (Bat SARSr-CoV HKU3-1)
- GenBank AID16716.1 (Bat SARSr-CoV Longquan-140)
- GenBank AVP78031.1 (Bat SARSr-CoV ZC45)
- GenBank AVP78042.1 (Bat SARSr-CoV ZXC21)
- GenBank QHR63300.2 (Bat CoV RaTG13)
- NCBI Reference Sequence YP_003858584.1 (Bat SARSr-CoV BM48-31)
- GISAID EPI_ISL_410721 (Pangolin BetaCoV Guandong2019)

Multiple sequence alignment of the RBD sequences was performed by MUSCLE version 3.8.31 [60]. Sequence logos were generated by WebLogo [61]. The conservation score of each RBD residue was calculated and mapped onto the SARS-CoV-2 RBD x-ray structure with ConSurf [62].

## ACKNOWLEDGEMENTS

We thank Henry Tien for technical support with the crystallization robot, Jeanne Matteson and Yuanzi Hua for contribution to mammalian cell culture, Wenli Yu for insect cell culture, Robyn Stanfield for assistance in data collection, and Paul Bieniasz for cells and plasmids for to the pseudovirus neutralization assays. We are grateful to the staff of Stanford Synchrotron Radiation Laboratory (SSRL) Beamline 12-1 for assistance. This work was supported by NIH K99 AI139445 (N.C.W.), the Bill and Melinda Gates Foundation OPP1170236 (A.B.W., I.A.W.), OPP1132237 and INV-002022 (R.W.S.) NIH HIVRAD P01 AI110657 (R.W.S., A.B.W., I.A.W.) and NIH CHAVD UM1 AI44462 (A.B.W., I.A.W.), the Netherlands Organization for Scientific Research (NWO) Vici grant (R.W.S.), the Fondation Dormeur, Vaduz (M.J.v.G.), a Health Holland PPS-allowance LSHM20040 (M.J.v.G.). M.J.v.G. is a recipient of an AMC Fellowship and a COVID-19 grant of the Amsterdam Institute of Infection and Immunity. J.v.S. is a recipient of a 2017 AMC Ph.D. Scholarship. Use of the SSRL, SLAC National Accelerator Laboratory, is supported by the U.S. Department of Energy, Office of Science, Office of Basic Energy Sciences under Contract No. DE-AC02-76SF00515. The SSRL Structural Molecular Biology Program is supported by the DOE Office of Biological and Environmental Research, and by the National Institutes of Health, National Institute of General Medical Sciences (including P41GM103393).

## AUTHOR CONTRIBUTIONS

H.L., N.C.W., M.Y. and I.A.W. conceived and designed the study. H.L., N.C.W., M.Y., and C.C.D.L. expressed and purified the proteins for crystallization. T.G.C., P.J.M.B., M.J.v.G. and R.W.S. provided antibody clones and sequences. T.G.C. performed binding analyses and J.v.S. provided neutralization data. H.L., N.C.W., M.Y. and X.Z. crystallized and determined the X-ray structures. S.B., J.L.T., and A.B.W. provided nsEM data and reconstructions. H.L., N.C.W., M.Y., and I.A.W. wrote the paper and all authors reviewed and/or edited the paper.

## COMPETING INTERESTS

Amsterdam UMC previously filed a patent application on the SARS-CoV-2 antibody COVA1-16 described here [6].

## DATA AVAILABILITY

X-ray coordinates and structure factors are being deposited to the RCSB Protein Data Bank. The COVA1-16 IGVH and IGVK sequences are available in GenBank: MT599919 and MT599835. The plasmids encoding the COVA1-16 IgG and Fab will be available from M.J.v.G. and R.W.S. under an MTA with the Amsterdam UMC. Other materials related to this paper will be available on request from the corresponding author.

---

**Supplementary Figure 1.**
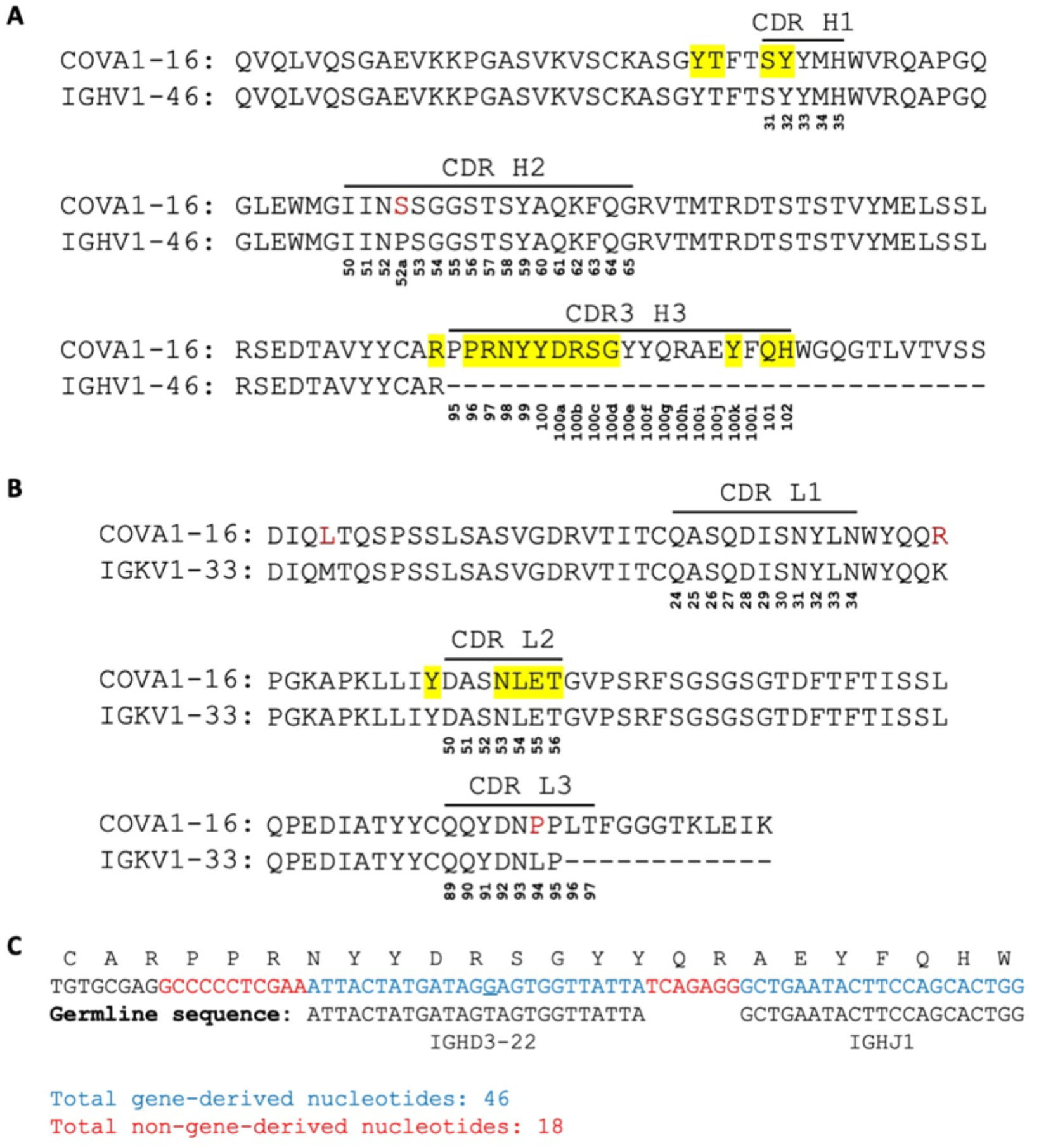
Comparison of COVA1-16 and putative germline sequences. Alignment of COVA1-16 Fab amino-acid sequence with **(A)** germline IGHV1-46 sequence, and **(B)** germline IGKV1-33 sequence. The regions that correspond to CDR H1, H2, H3, L1, L2, and L3 are indicated. Residues that differ from germline are highlighted in red. COVA1-16 Fab residues that interact with the RBD are highlighted in yellow. Residue positions in the CDRs are labeled according to the Kabat numbering scheme. **(C)** Amino acid and nucleotide sequences of the V-D-J junction of COVA1-16, with putative gene segments (blue) and N-regions (red), are indicate. The germline sequences of IGHD3-22 and IGHJ1 are also shown. The only somatically mutated nucleotide in the D region is underlined.

**Supplementary Figure 2.**
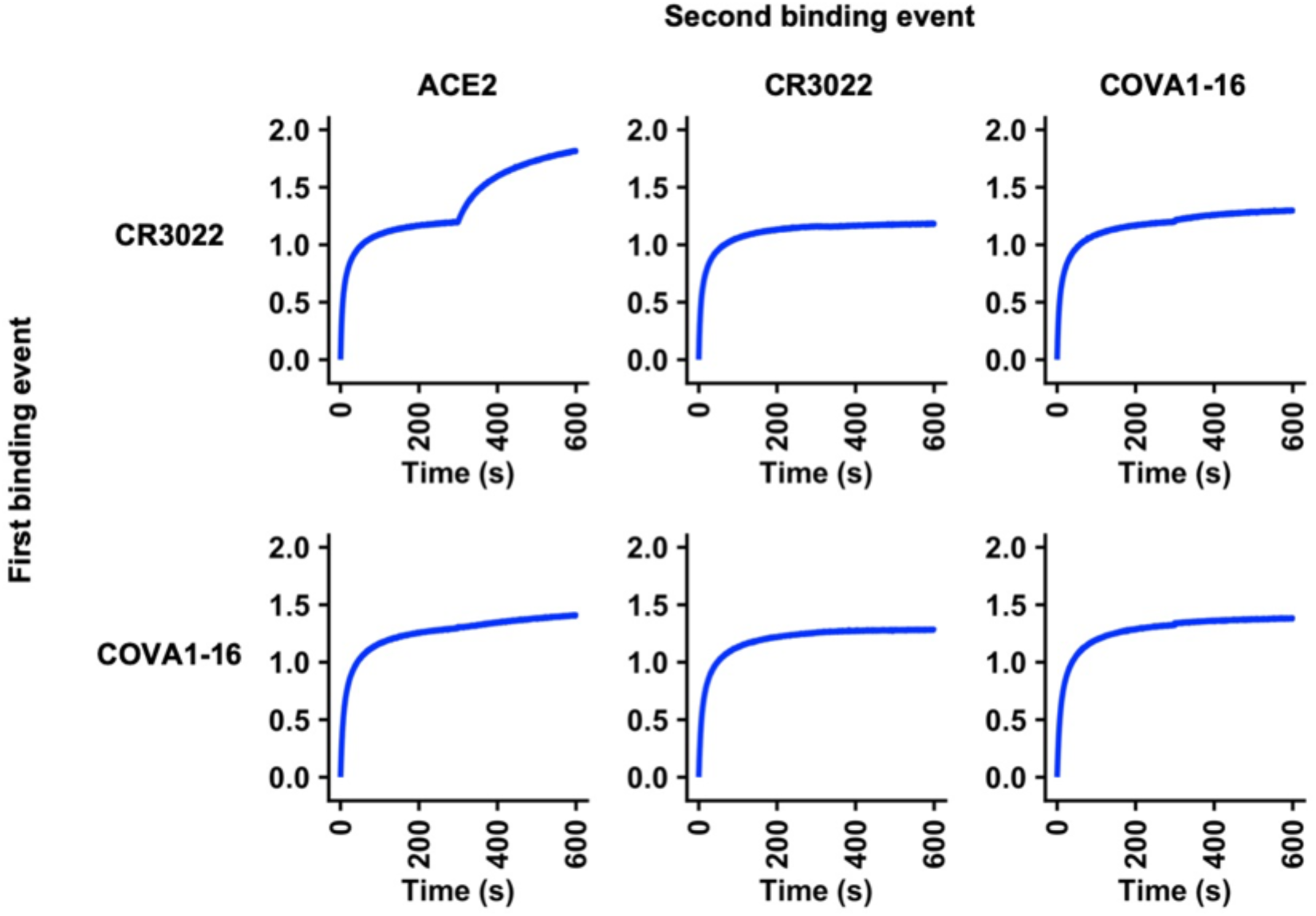
Competition assay between different IgGs and ACE2. Competition between COVA1-16 IgG, CR3022 IgG, and Fc-tagged ACE2 was measured by biolayer interferometry (BLI). Y-axis represents the response. The biosensor was first loaded with SARS-CoV-2 RBD, followed by two binding events: 1) CR3022 IgG or COVA1-16 IgG, and 2) ACE2, CR3022 IgG, or COVA1-16 IgG. A period of 300 s was used for each binding event. A further increase in signal during the second binding event (starting at 300 s time point) indicates lack of competition with the first ligand.

**Supplementary Figure 3.**
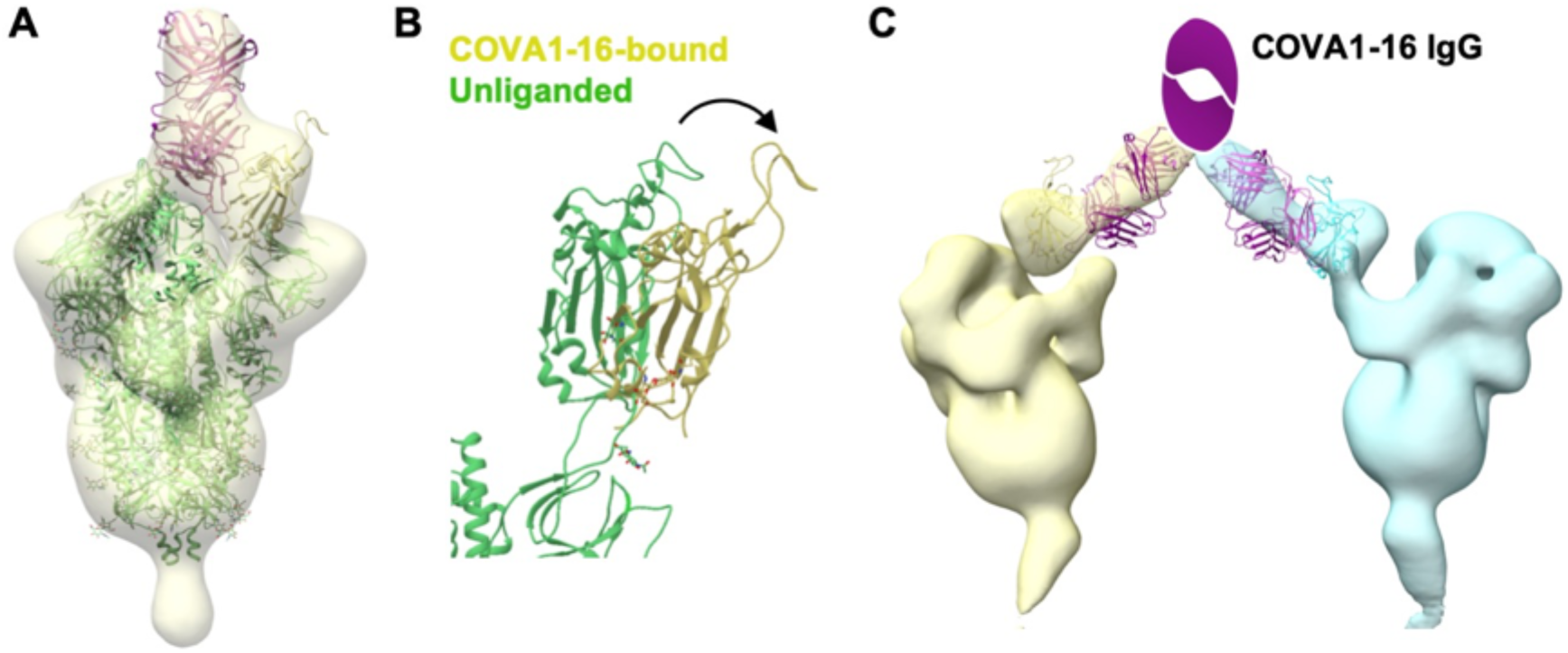
Negative-stain EM analysis of COVA1-16 binding to SARS-CoV-2 S trimer. **(A)** An atomic model from the crystal structure of SARS-CoV-2 RBD bound to COVA1-16 Fab was fit into the negative-stain EM reconstruction of the SARS-CoV-2 spike bound to COVA1-16 Fab. The COVA1-16 Fab approaches the apex of the S trimer in a perpendicular orientation. A secondary structure backbone representation of the prefusion spike model (PDB: 6Z97, green) [1] was also fit into the EM density with RBD residues (334-528) removed from one of the protomers here for clarity. The COVA1-16 heavy and light chains are in magenta and pink, respectively, and COVA1-16-bound RBD in yellow. **(B)** Conformation of RBD in an up conformation from an unliganded SARS-CoV-2 S trimer (PDB: 6Z97, green) [1] is compared to that of the RBD (yellow) bound by COVA1-16 Fab. The arrow indicates that the RBD further rotates and opens up when bound to COVA1-16, thereby moving further away from the trimer threefold axis. **(C)** An atomic model of the spike RBD bound to COVA1-16 Fab is fit into a negative-stain EM reconstruction, where COVA1-16 Fab approaches the SARS-CoV-2 S trimer from the side. COVA1-16 is modelled as an IgG to illustrate the feasibility of bivalent binding to adjacent spike proteins on the virus surface. The Fab heavy and light chains are shown in magenta and pink. A schematic representation of the Fc domain of the IgG is shown in magenta. The RBD model and spike density for each trimer is shown in yellow and cyan.

**Supplementary Figure 4.**
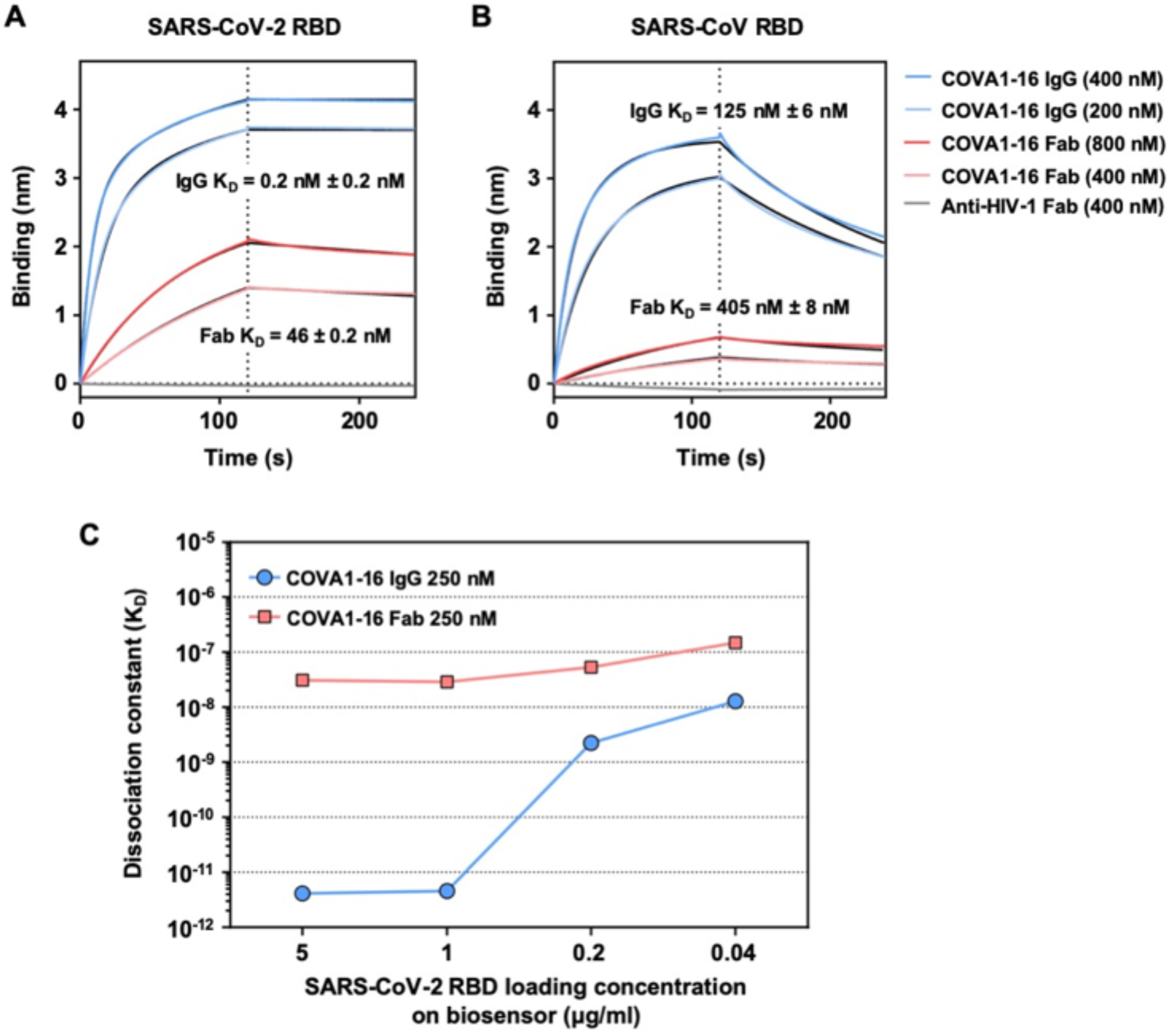
Sensorgrams for binding of COVA1-16 to SARS-CoV-2 RBD and SARS-CoV RBD. **(A-B)** Binding kinetics of COVA1-16 Fab and IgG to **(A)** SARS-CoV-2 RBD and **(B)** SARS-CoV RBD were measured by biolayer interferometry (BLI) with RBD on the biosensor and antibody in solution. Y-axis represents the response. An anti-HIV His-tagged Fab (4E1) was used as a negative control. Dissociation constants (K_D_) for IgG and Fab were obtained using a 1:2 bivalent model and 1:1 binding model, respectively, which are represented by the black lines. Representative results of two replicates for each experiment are shown. **(C)** The relationship between SARS-CoV-2 RBD loading concentration on the biosensor and the dissociation constant of COVA1-16 IgG is shown.

**Supplementary Figure 5.**
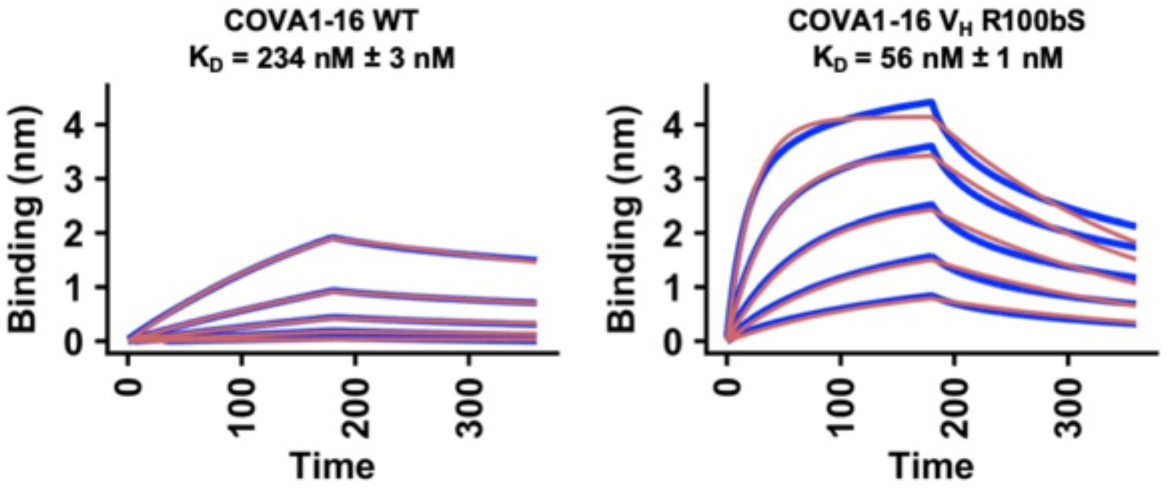
Sensorgrams for binding of COVA1-16 wild-type and V_H_ R100bS mutant Fabs to SARS-CoV-2 RBD. Binding kinetics of COVA1-16 wild-type and V_H_ R100bS mutant Fab to SARS-CoV-2 RBD were measured by biolayer interferometry (BLI) with RBD on the biosensor and antibody in solution. Y-axis represents the response. Dissociation constants (K_D_) for Fabs were obtained using a 1:1 binding model, which are represented by the red lines. Representative results of two replicates for each experiment are shown. Unlike Figure S4, which used HEK293F-expressed SARS-CoV-2, the experiment here used insect cell-expressed SARS-CoV-2.

**Supplementary Figure 6.**
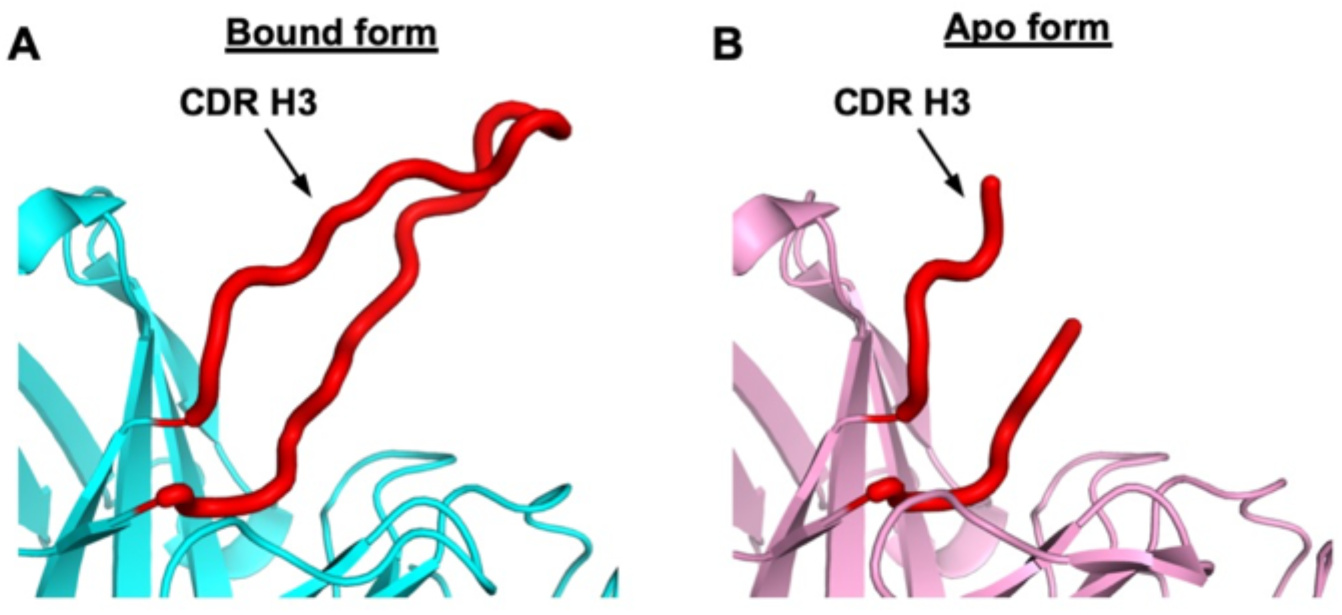
CDR H3 of COVA1-16 Fab is disordered in its unliganded apo form. **(A)** In the crystal structure of the RBD-bound form of COVA1-16 Fab, the CDR H3 loop is completely ordered (red). **(B)** In the crystal structure of the apo form of COVA1-16, the distal end of the CDR H3 loop is intrinsically disordered or flexible (red).

**Supplementary Figure 7.**
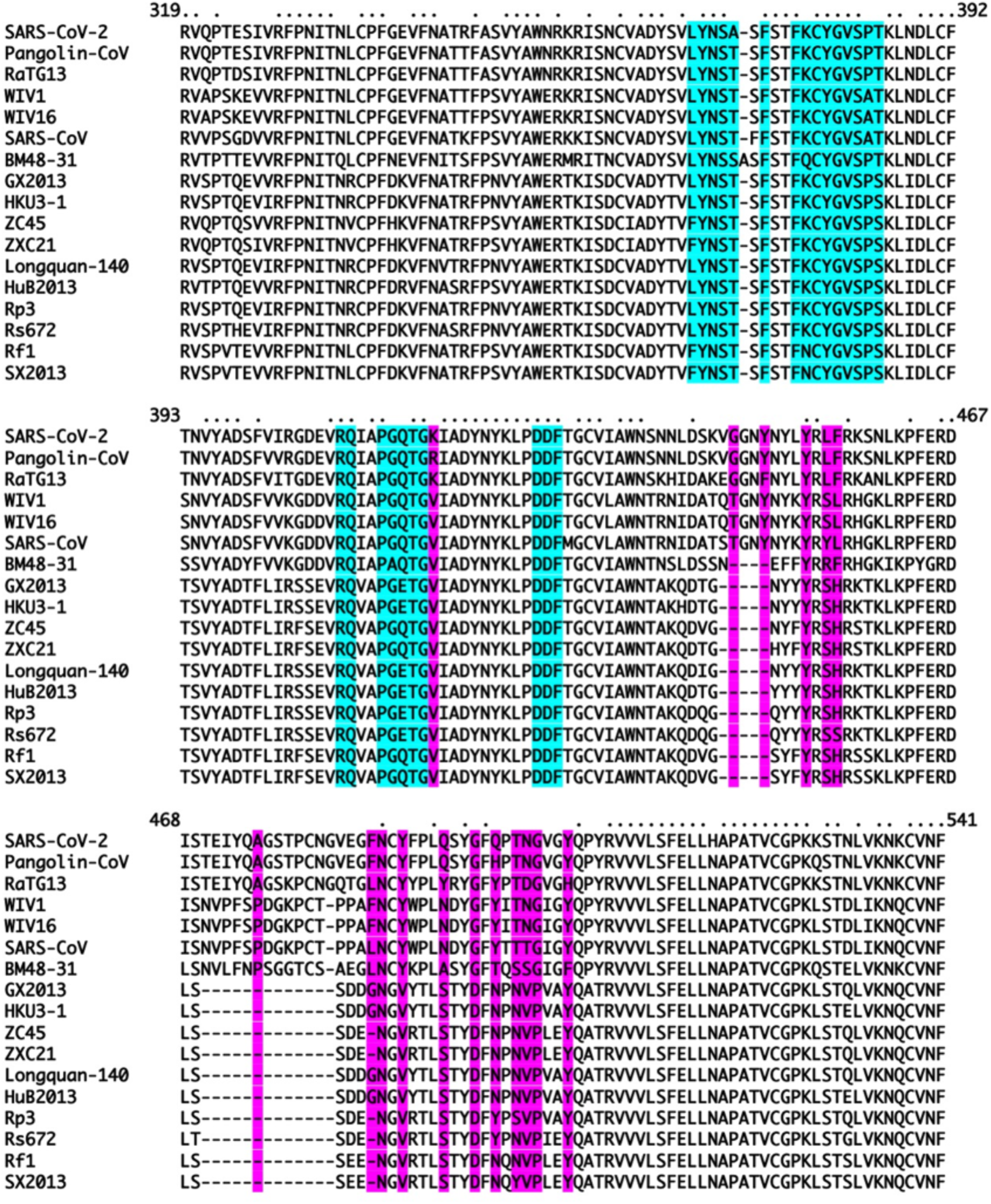
Sequence alignment of the RBD from SARS-related coronaviruses. Amino-acid sequences of RBDs from SARS-CoV-2, SARS-CoV, and other SARS-related coronavirus (SARSr-CoV) strains are aligned. COVA1-16 epitope residues are highlighted in cyan. ACE2-binding residues are highlighted in purple. Conserved residues are indicated by small black dots on the top of the alignment.

**Supplementary Figure 8.**
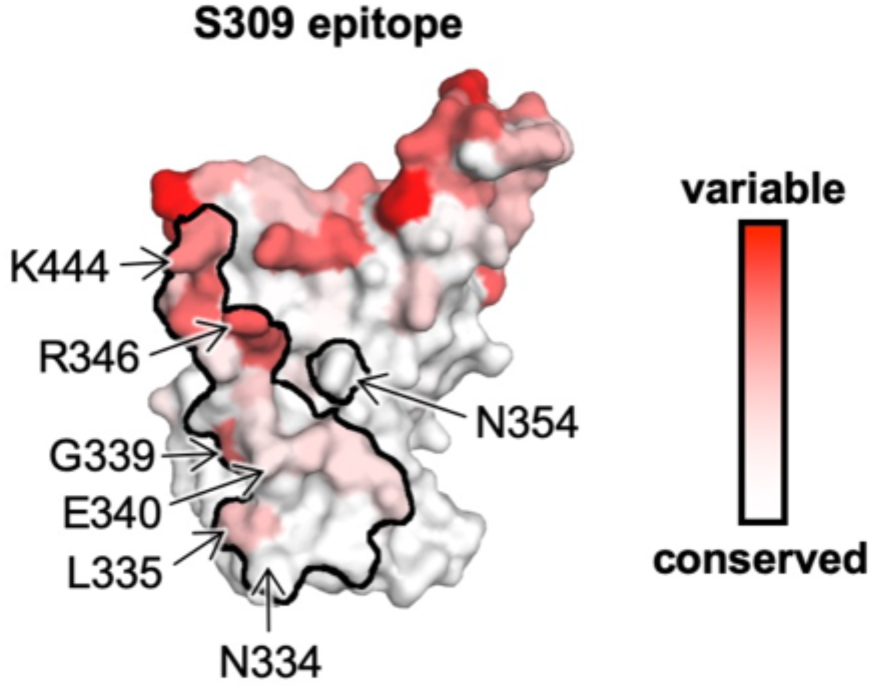
Sequence conservation of S309 epitope. Sequence conservation of the RBD is highlighted on the structure for S309 epitope [2]. This view corresponds to the opposite side (rotated 180 degrees along the vertical axis) from that shown in Figure 4A–B.

**Supplementary Figure 9.**
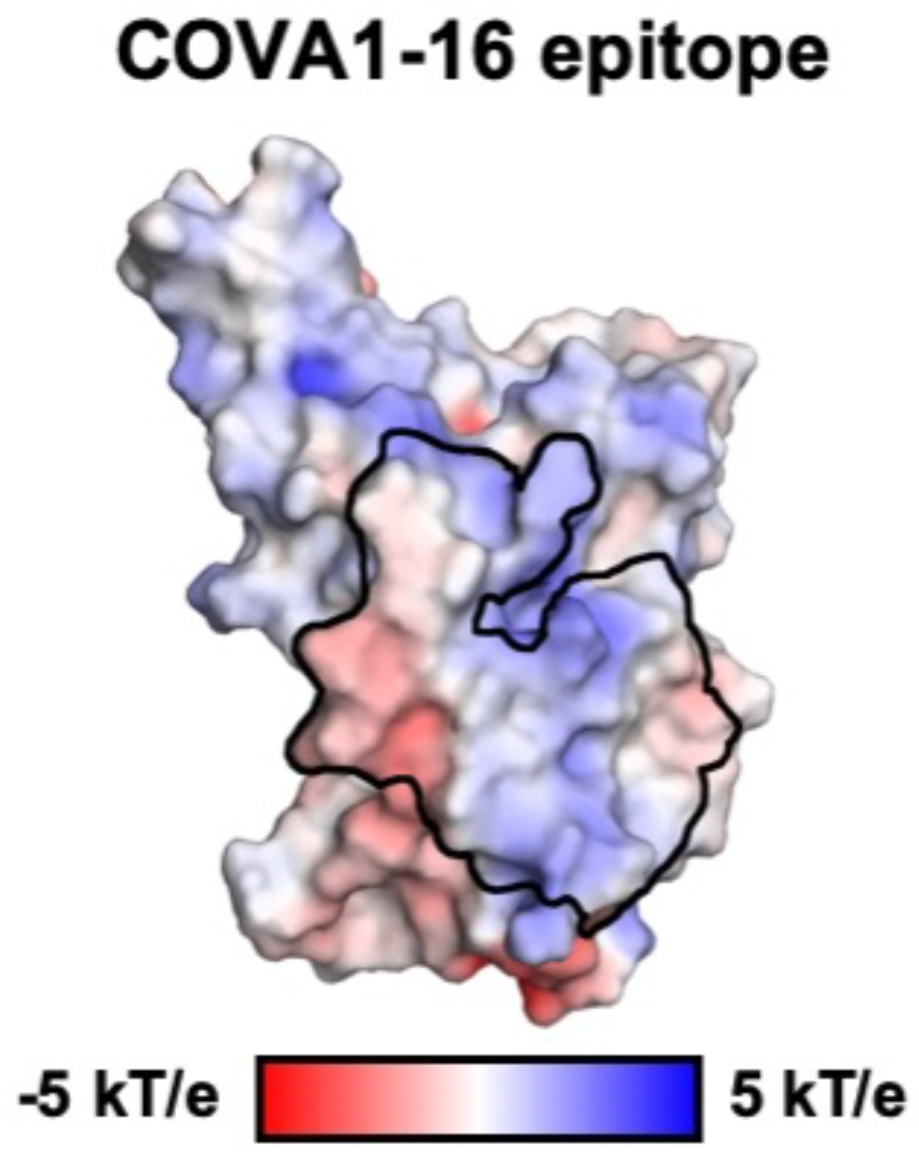
COVA1-16 epitope in electrostatic surface representation. The epitope of COVA1-16 is outlined and shows its largely polar nature.

**Supplementary Figure 10.**
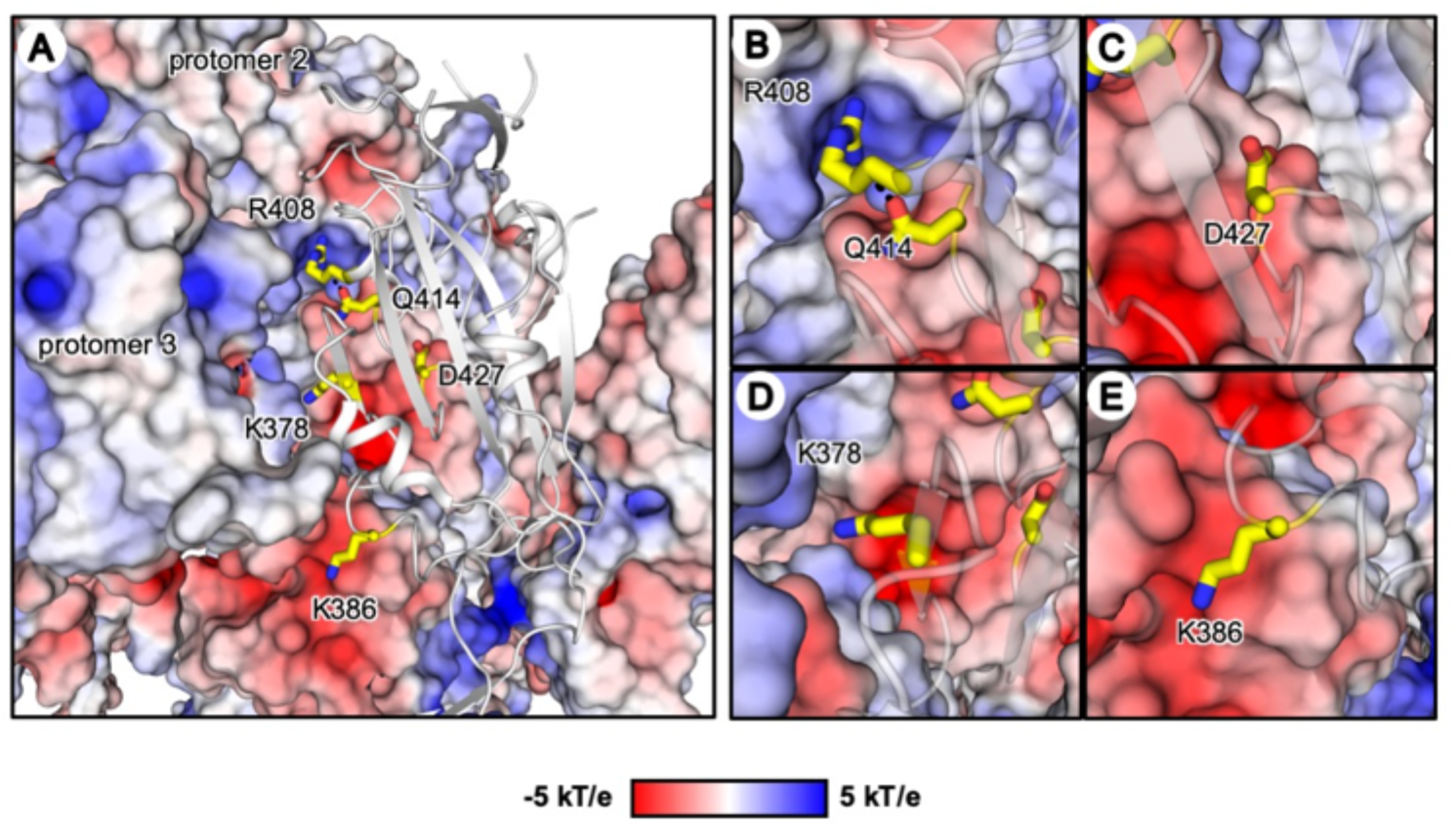
Location of residues of interest in the COVA1-16 epitope when all three RBDs are in the “down” conformation. **(A)** The RBD of one of the three protomers is shown as a gray cartoon with the side chains of five residues of interest shown in yellow stick representation. RBD residues K378, R408, Q414, and D427 are within the COVA1-16 epitope, whereas K386 is not a COVA1-16 epitope residue. The other two protomers (protomers 2 and 3) are shown in a surface electrostatic representation. **(B-E)** Zoomed-in views for the regions surrounding residues **(B)** R408 and Q414, **(C)** D427, **(D)** K378, and **(E)** K386. A hydrogen bond in **(B)** is represented by a dashed line. Due to charge difference or similarity between the side chain and the proximal region of the neighboring protomer, either repulsive (same charge) or attractive (opposite charge) environments are found and visualized here. PDB 6VXX is used to represent the spike protein [3]. Of note, the shape complementarity values (Sc) [4] of the COVA1-16 epitope/RBD interface, COVA1-16 epitope/S2 interface, and COVA1-16 epitope/COVA1-16 interface are 0.53, 0.75, and 0.74, respectively, indicating good complementary and tight fit of the COVA1-16 epitope surface with the rest of the trimer in the RBD down conformation. Sc values can range from 0 to 1, with a larger Sc value represents higher shape complementarity.

**Supplementary Figure 11.**
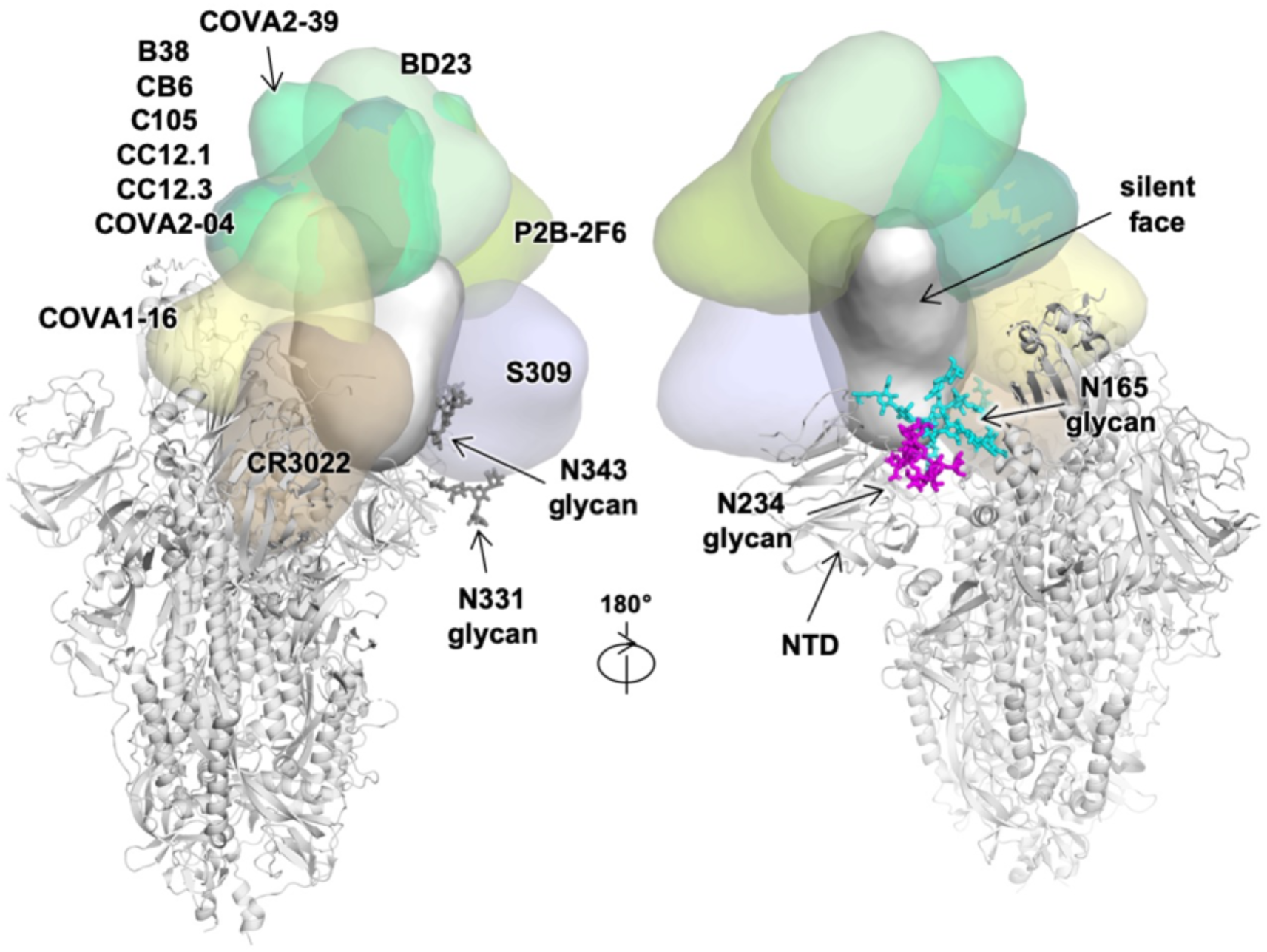
The N-glycan on the N-terminal domain (NTD) also shields part of the RBD. The antibody-bound RBD, which is displayed and colored as in Figure 5, is shown in the up conformation on the S protein (PDB 6VSB) [5]. N-glycans on N165 (NTD), N234, N331, and N343 (RBD) are modelled according to the main glycoform observed at these sites in [6], and shown in stick representation. Antibody Fabs from published crystal and cryo-EM structures are represented as globular outlines in different colors as outlined in Figure 5. B38, CB6, C105, CC12.1, CC12.3, COVA2-04, COVA2-39, BD23, P2B-2F6 all bind at or around the receptor binding site. S309 binds to the elongated accessible face of the RBD in both up and down conformations, and CR3022 binds to the opposite face that is exposed in the RBD up conformation, but buried in the RBD down conformation.

**Supplementary Figure 12.**
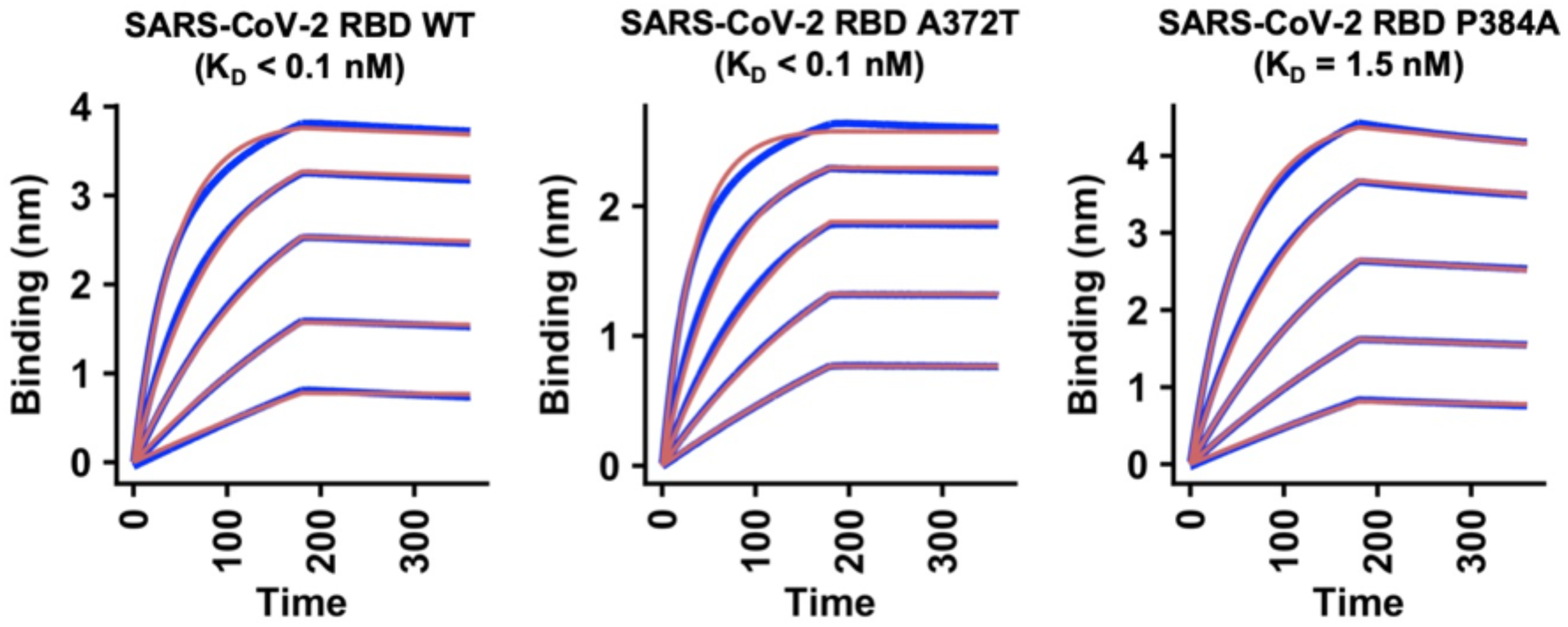
Sensorgrams for binding of COVA1-16 IgG to SARS-CoV-2 RBD WT or mutants. Binding kinetics of COVA1-16 IgG to SARS-CoV-2 RBD WT, A372T, and P384A were measured by biolayer interferometry (BLI) with RBD on the biosensor and antibody in solution. Y-axis represents the response. Dissociation constants (K_D_) for Fabs were obtained using a 1:1 binding model, which are represented by the red lines. Representative results of two replicates for each experiment are shown. A372T and P384A are the only two mutations that differ between the SARS-CoV-2 and SARS-CoV sequences in COVA1-16 epitope. The affinity of COVA1-16 IgG to the A372T mutant did not show any detectable difference from WT. Although the affinity (K_D_) of COVA1-16 IgG to the P384A mutant decreases, the binding is still 100 times tighter than that measured between COVA1-16 IgG and SARS-CoV RBD (Figure S4B). As a result, the binding affinity of COVA1-16 to the RBD may be influenced by residues outside of the epitope as well as the dynamics of the RBD fluctuations between up and down conformations.

**Supplementary Table 1.**
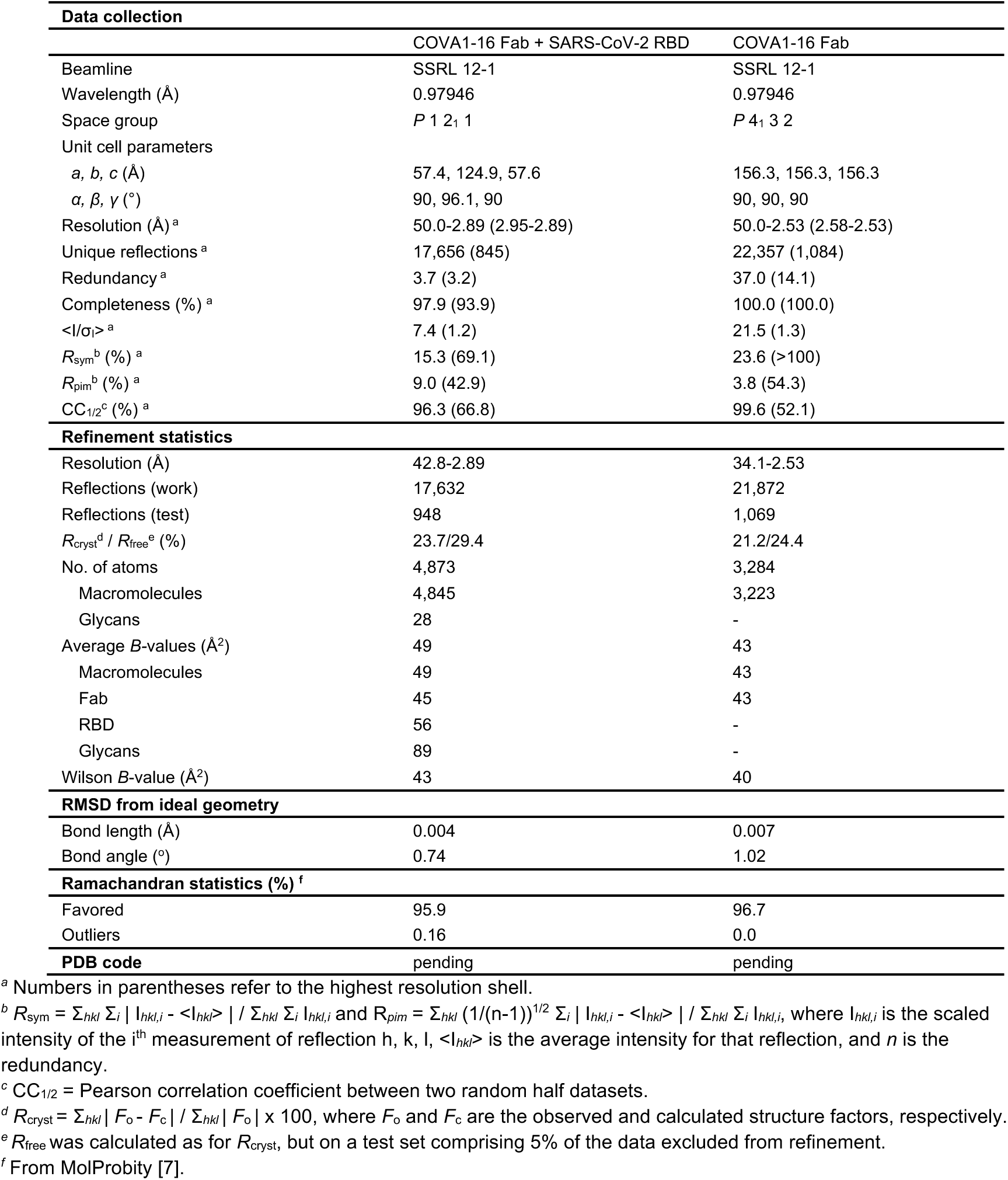
X-ray data collection and refinement statistics

**Supplementary Table 2.** Hydrogen bonds identified in the antibody-RBD interface using the PISA program

| **COVA1-16 Fab** | **Distance**<br>**[Å]** | **SARS-CoV-2 RBD** |
| --- | --- | --- |
| H:ARG100b[NH2] | 3.3 | A:TYR369[O] |
| H:ARG100b[NE] | 3.9 | A:SER371[O] |
| H:ARG100b[N] | 3.8 | A:PHE377[O] |
| H:TYR100[N] | 2.6 | A:CYS379[O] |
| H:GLN101[NE2] | 3.1 | A:GLN414[OE1] |
| H:ARG97[NH1] | 2.5 | A:ASP427[O] |
| H:TYR32[OH] | 3.1 | A:ASP427[OD1] |
| H:THR28[ N] | 3.2 | A:ASP427[OD2] |
| H:ARG97[NH1] | 3.0 | A:PHE429[O] |
| H:TYR100[O] | 2.9 | A:CYS379[N] |
| H:SER100c[O] | 3.3 | A:THR385[OG1] |
| H:GLN101[OE1] | 3.8 | A:GLN414[NE2] |
| L:ASN53[OD1] | 3.2 | A:ARG408[NH2] |
| L:LEU54[O] | 3.7 | A:ARG408[NE] |

